# Evaluating the Robustness of Connectivity Methods to Noise for In Silico Drug Repurposing Studies

**DOI:** 10.1101/2022.09.14.508054

**Authors:** Nevin Tham, Sarah R Langley

## Abstract

Drug repurposing is an approach to identify new therapeutic applications for existing drugs and small molecules. It is a field of growing research interest due to its time and cost effectiveness as compared with *de novo* drug discovery. One method for drug repurposing is to adopt a systems biology approach to associate molecular ‘signatures’ of drug and disease. Drugs which have an inverse relationship with the disease signature may be able to reverse the molecular effects of the disease and thus be candidates for repurposing. Conversely, drugs which mimic the disease signatures can inform on potential molecular mechanisms of disease. The relationship between these disease and drug signatures are quantified through connectivity scores. Identifying a suitable drug-disease scoring method is key for in silico drug repurposing, so as to obtain an accurate representation of the true drug-disease relationship. There are several methods to calculate these connectivity scores, notably the Kolmogorov-Smirnov (KS), Zhang and eXtreme Sum (XSum). However, these methods can provide discordant estimations of the drug-disease relationship and this discordance can affect the drug-disease indication. Using the gene expression profiles from the Library of Integrated Network-Based Cellular Signatures (LINCS) database, we evaluated the methods based on their drug-disease connectivity scoring performance. In this first-of-its-kind analysis, we varied the quality of disease signatures by using only highly differential genes or by the inclusion of non-differential genes. Further, we simulated noisy disease signatures by introducing varying levels of noise into the gene expression signatures. Overall, we found that there was not one method that outperformed the others in all instances, but the Zhang method performs well in a majority of our analyses. Our results provide a framework to evaluate connectivity scoring methods, and considerations for deciding which scoring method to apply in future systems biology studies for drug repurposing.

## 1 Introduction

Drug repurposing is the process of identifying a new therapeutic use for an existing drug. It is a field of growing research interest because the traditional process to discover and develop novel therapeutics is long and expensive with low rates of success. As the pharmacokinetics and toxicology of approved drugs have been well studied, it renders drug repurposing markedly more economical and efficient (Breckenridge and Jacob 2019; Pushpakom et al. 2019). To this end, the Library of Integrated Network-Based Cellular Signatures (LINCS) Project was conceived to facilitate in silico drug repurposing (Subramanian et al. 2017). It involves the generation of a large-scale compendium of transcriptional profiles obtained from small molecule compound perturbations in human cultured cell lines. Over the years, there have been several reported applications on the use of LINCS data for in silico drug repurposing in diseases such as epilepsy (Mirza et al. 2017), diabetes (Jin et al. 2014) and cancer (Rho, Kim, and Kang 2011; Lim, Lim, and Cho 2014; H.-W. Cheng et al. 2015).

The concept behind in silico drug repurposing is to identify connections between the transcriptional profile of a disease and small molecule compounds (Lamb et al. 2006). The mRNA expression state of a cell captures information about the transcriptional regulation processes taking place in the cell. Hence, by comparing the fingerprints of gene expression induced by a drug and a disease, the association between the drug and the disease can be evaluated. To quantify the relationship between a drug and a disease, the direction and magnitude of gene expression changes in the disease signature are compared against that in the drug signature. A drug that increases the gene expression of down-regulated genes (and decreases gene expression of up-regulated genes) in the disease signature is defined to have an inverse drug-disease relationship and is predicted to be able to reverse the disease signature. On the other hand, a drug that increases the gene expression of up-regulated genes (and decreases gene expression of down-regulated genes) in the disease signature, has a positive drug-disease relationship and is predicted to reflect or phenocopy the disease. Using similarity scores (or connectivity scores) between the disease signature and drug signatures, such as those found in LINCS, it is possible to prioritise drugs that can be investigated as potential treatments for the disease.

There have been several proposed algorithms to quantify the similarity between two transcriptional signatures. The first algorithm was adopted by Lamb et al., and it uses a nonparametric rank-based algorithm, based on the Kolmogorov-Smirnov (KS) statistic (Lamb et al. 2006). This method was also later referred to as the Gene Set Enrichment Analysis (GSEA) method. Zhang and Gant next introduced a simpler method, known as the Zhang (or statistically significant connectivity map – ssCMap) method (Zhang and Gant 2008). Unlike the KS method, the Zhang method takes account of the direction of regulation of the genes in the reference profile, and it is based on the signed-rank statistic. A third method, eXtreme sum (XSum), proposes that a reference profile can be represented by the most highly up and down regulated genes, known as eXtreme genes (J. Cheng et al. 2014). The fold changes of these genes are then used to quantify the connection between the two signatures. Besides the XSum algorithm, Cheng and Yang explored several other pairwise similarity metrics which utilises eXtreme genes, such as XCosine, XCorrelation and XSpearman. Among the eXtreme methods, the XSum method is recommended due to its ease of use and minimal information required (J. Cheng and Yang 2013).

Given multiple gene-based connectivity algorithms, it has been of interest to benchmark the methods. The common approach to evaluate the methods is to first compute drug-drug similarity, and then assess the true relationship of the compounds using their Anatomical Therapeutic Chemical (ATC) classification (Iorio, Tagliaferri, and Bernardo 2009; Iskar et al. 2010). However, little work has been done to evaluate the methods based on their ability to quantify drug-disease dissimilarity. It has been highlighted that the validity of such studies may be limited as the quality of the disease signature may not completely portray the disease profile (Musa et al. 2018). Moreover, the influence of the disease signature quality on connectivity score is an aspect of in silico drug repurposing that is widely recognised, yet relatively understudied (Zhang and Gant 2008; J. Cheng et al. 2014; Musa et al. 2018).

Hence in this work, we evaluated three primary drug repurposing methods — KS, Zhang, XSum — based on their ability to identify approved, or experimentally validated, drugs for repurposing to treat other diseases. The disease signatures were queried against the most recent release of the LINCS database, which contains over a million replicate-collapsed signatures. In addition to benchmarking against the drug-drug signatures that have previously been evaluated, we take a first step to investigate the drug-disease indications of these methods when the quality of the disease signature is varied. We further explored how compound prioritisation by these methods are altered due to noise in gene expressions. We show that the Zhang method generally had a better sensitivity, and is more robust to variation in query signature quality, than the other two methods. Together, this suggests that the Zhang method is better suited to quantify drug-disease relationships for in silico drug repurposing.

## 2 Methods

### 2.1 Query Signatures from Literature

To evaluate the relative performance of different scoring algorithms, four query signatures were obtained from literature. They comprise three disease signatures and one drug signature. Table 1 summarises these publicly available query signatures. As the epilepsy signature was obtained from the mouse model, the mouse genes were mapped to their analogous human genes to perform the analyses.

**Table 1:** Drug and disease signatures used to query LINCS drug signature database.

| Query Signature | Type | Platform | Organism | Source |
| --- | --- | --- | --- | --- |
| 17b-estradiol (E2) | Drug | Microarray | Human | Frasor et al. (2004) |
| Gastric Cancer | Disease | Microarray | Human | Claerhout et al. (2011) |
| Colorectal Cancer | Disease | RNA-Seq | Human | Noort et al. (2014) |
| Epilepsy | Disease | RNA-Seq | Mouse | Hansen et al. (2014) |

### 2.2 LINCS Database

The LINCS level 5 data of compound-treated signatures were downloaded from Clue.io (https://clue.io/data/CMap2020#LINCS2020 (CMap LINCS 2020)). Level 5 data in CMap refers to replicate consensus signatures, where the differential expression (LFC) of genes are moderated across replicated experiments to determine the ‘de-noised’ representation of a drug effect (Subramanian et al. 2017). This database comprises more than 720K signatures of 34K compounds treated onto 248 unique cell lines. The drug signature database was further filtered down to signatures which were only treated with compounds with known targets and MOAs, resulting in 387K signatures of 2,558 unique compounds.

In order to incorporate noise into the LINCS data, the level 3 data was also retrieved from the same web source as the level 5 data. LINCS level 3 data refers to individual instances of normalised gene expressions, due to a functional perturbation. The level 3 data which was retrieved comprises 1.8M compound treated instances.

### 2.3 Drug-Disease/Drug-Drug Similarity Score

The three in silico drug repurposing methods discussed in this study are the KS, Zhang and XSum methods. Since the introduction of the concept of drug-disease scoring, there have been several other methods proposed to quantify the relationship between a drug and a disease signature. Most of these other methods are a modification or a weighted combination of these three parent methods. Hence, this study focuses on the evaluation of these three primary methods. The R code used for these three methods was retrieved from the RCSM package, which was compiled in an earlier work (Lin et al. 2020). A brief description of their algorithm is summarised in Table 2.

**Table 2:** Brief description of current drug-disease algorithms that are evaluated in this study.

| Method | Description | Reference |
| --- | --- | --- |
| Kolmogorov-Smirnov (KS) | First proposed method to quantify drug-disease relationship.<br>Genes in the drug and disease signatures are ranked by their LFC.<br>Determine the KS statistic for up and down regulated genes, based on the maximum difference between the relative ranks of genes in drug and disease signatures.<br>Compute difference between the KS statistic of the up and down regulated genes. | Lamb et al. (2006) |
| Zhang | Genes are ranked based on their absolute LFC, where genes with largest absolute LFC are assigned with the largest numerical rank.<br>Direction of gene regulation in the drug signature (+1 for up-regulated, -1 for down regulated) is multiplied with the numerical rank.<br>Sum product of actual gene ranks is divided against the maximum theoretical sum product of gene ranks. | Zhang and Gant (2008) |
| eXtreme Sum (XSum) | Subsets the top and bottom $N$ DEGs by LFC in the drug signature.<br>Intersect the above genes with DEGs of the disease (eXtreme genes).<br>Determine the sums of drug LFC of eXtreme genes. | J. Cheng et al. (2014) |

#### 2.3.1 Kolmogorov-Smirnov Statistic

The benchmark algorithm proposed by (Lamb et al. 2006) uses a non-parametric, rank-based pattern-matching strategy based on the Kolmogorov-Smirnov (KS) statistic. Essentially, this method calculates the maximum difference in relative ranks of the up and down regulated genes. A strongly down-regulated gene by the disease that is strongly up-regulated by the drug, results in a large difference between their relative rank. This contributes to a strongly negative drug-disease score, indicating the strong potential of the drug to reverse the disease signature.

#### 2.3.2 Zhang Score

Zhang et al. proposed the Zhang scoring algorithm, and have shown that their method performs better than the KS method (Zhang and Gant 2008). The Zhang Score (ZS) is obtained by finding the ratio of the sum product of the actual gene ranks against the maximum theoretical sum product of the gene ranks. The Zhang method ranks the gene importance based on the absolute value of the gene log fold change (LFC), and assigns heavier weights to the more differentiated genes. It is a sign-rank based algorithm, which assigns a negative rank for genes that are down-regulated, and vice versa. They rationalise that ZS is a more accurate quantification of the drug-disease relationship because the highly regulated genes contribute more to the ZS, and are assigned greater weights regardless of the direction of regulation.

#### 2.3.3 eXtreme Sum

Cheng et al. proposed the eXtreme Sum (XSum) scoring method, which is based on eXtreme genes (J. Cheng et al. 2014). eXtreme genes are genes that have been changed by the disease and also within the top and bottom *N* genes changed by the drug (*N* is an arbitrary integer defined by the user). XSum can be computed by: (sum of the LFC of the up-regulated eXtreme genes) – (sum of the LFC of the down-regulated eXtreme genes). As the size of baseline query signatures used in this study ranges between 80 to 120 genes, *N* is set to 150 for this analysis, so that the number of eXtreme genes may be comparable with the query signature. The XSum is also a sign-rank based algorithm, wherein the signs and ranks of the genes influence the overall score.

### 2.4 Comparison by Number of Significantly Scored Signatures

Complementary to calculating drug-disease scores, the significance of a score is also evaluated. This is achieved by comparing the actual drug-disease score against a null distribution of scores. The null distribution of scores is obtained by computing scores between a drug signature and multiple randomly generated disease signatures of identical size. The significance (or empirical p-value) of the actual drug-disease score is determined by the frequency at which its absolute value exceeds the absolute scores of the null distribution (Zhang and Gant 2008; Lin et al. 2020). For the case of a positive (negative) drug-disease score, the p-value indicates if the phenocopy (reversal) score is significantly positive (negative, p < 0.05), or positive (negative) by chance (p ≥ 0.05).

To compare which method was able to significantly score the highest number of true phenocopy (or reversal) signatures, the literature query signatures were queried against signatures that have been perturbed with a compound of the same MOA (or with an actual drug that has been used to treat the disease) (Table 3). The method that has the largest number of significantly scored phenocopy or reversal signatures suggests that it has a better sensitivity.

**Table 3:** List of drugs that are marked as phenocopy or reversal of the literature signature, in order to investigate the accuracy of each method. * compound not currently used to treat the disease, but experimental validation has shown its potential to reverse disease pathology. ESR: estrogen receptor.

| Query Signature | Direction | (Types of) Drugs |
| --- | --- | --- |
| 17b-estradiol | Phenocopy | ESR agonists |
| 17b-estradiol | Reversal | ESR antagonists |
| Colorectal Cancer | Reversal | Capecitabine, Fluorouracil, Irinotecan, Regorafenib, Citalopram*, Troglitazone* |
| Epilepsy | Reversal | Lamotrigine, Carbamazepine, Levetiracetam, Topiramate, Sitagliptin* |

For the purpose of this analysis, the signatures in LINCS that were treated with the corresponding drugs (Table 3) regardless of the dosage or treatment duration, are generalised as true phenocopy or reversal signatures. In truth, an excessive (or insufficient) dosage of a drug for an extended (or shortened) duration, may alter the status of a signature as a true positive. As such, these generalised signatures are only putative true positives, and it is unlikely that all of them are true positives. Consequently, the number of true positive signatures cannot be exactly determined, and the score obtained from this analysis is only a close estimate of the true accuracy.

### 2.5 Comparison by AUC of Signature Score

To evaluate the relative performance of each scoring metric using the area under the Receiver Operator Curve (ROC, AUC), the set of reversal and non-reversal drugs were first defined. Compounds that have been used to treat the disease were defined as reversal drugs (Table 3), while the complement set of compounds in LINCS were defined as drugs that do not reverse the disease. These compound labels were then used to compute the AUC.

The ROC records how sensitivity of a method changes with respect to its specificity; and methods that attain larger AUC indicate their better accuracy. In the application of drug repurposing, it is prudent to investigate only the top scoring drugs. Hence, the early retrieval performance is also evaluated, which is measured by the AUC at which the false positive rate (FPR) is less than 0.1 and 0.01 (AUC0.1 and AUC0.01) (J. Cheng et al. 2014).

### 2.6 Comparison by Varying Signature Quality

In the formulated methods to quantify drug-disease score, a common caveat is that the quality of the disease signature may affect the performance of the method. The quality of a disease signature can be estimated by the proportion of significant DEGs, as well as their significance level in the differential expression analysis. To evaluate the how performance of each method is affected by data quality, we simulated and tested varying qualities of the gastric cancer and epilepsy disease signatures across five different levels (Table 4). For each quality level, the drug-disease scores were calculated against a specific drug signature in LINCS (gastric cancer: ASG002_AGS_24H:G15, epilepsy: REP.A006_MCF7_24H:D05), which had obtained a significantly negative score with the baseline medium sized disease signature (set A). The medium sized signature set A was used as the baseline for this analysis, to understand the score trend when the query signature quality changes. Set A was arbitrarily derived from the top 250 and 120 genes by order statistics in the gastric cancer and epilepsy DEG analysis, respectively (order statistics: *abs*(*LFC*) × −*log*_10_(*adjusted p* − *value*)). Set B is derived from the top 120 and 60 genes, while Set C is derived the top 500 and 250 genes, by order statistics in the gastric cancer and epilepsy DEG analysis, respectively (Table 4).

**Table 4:**
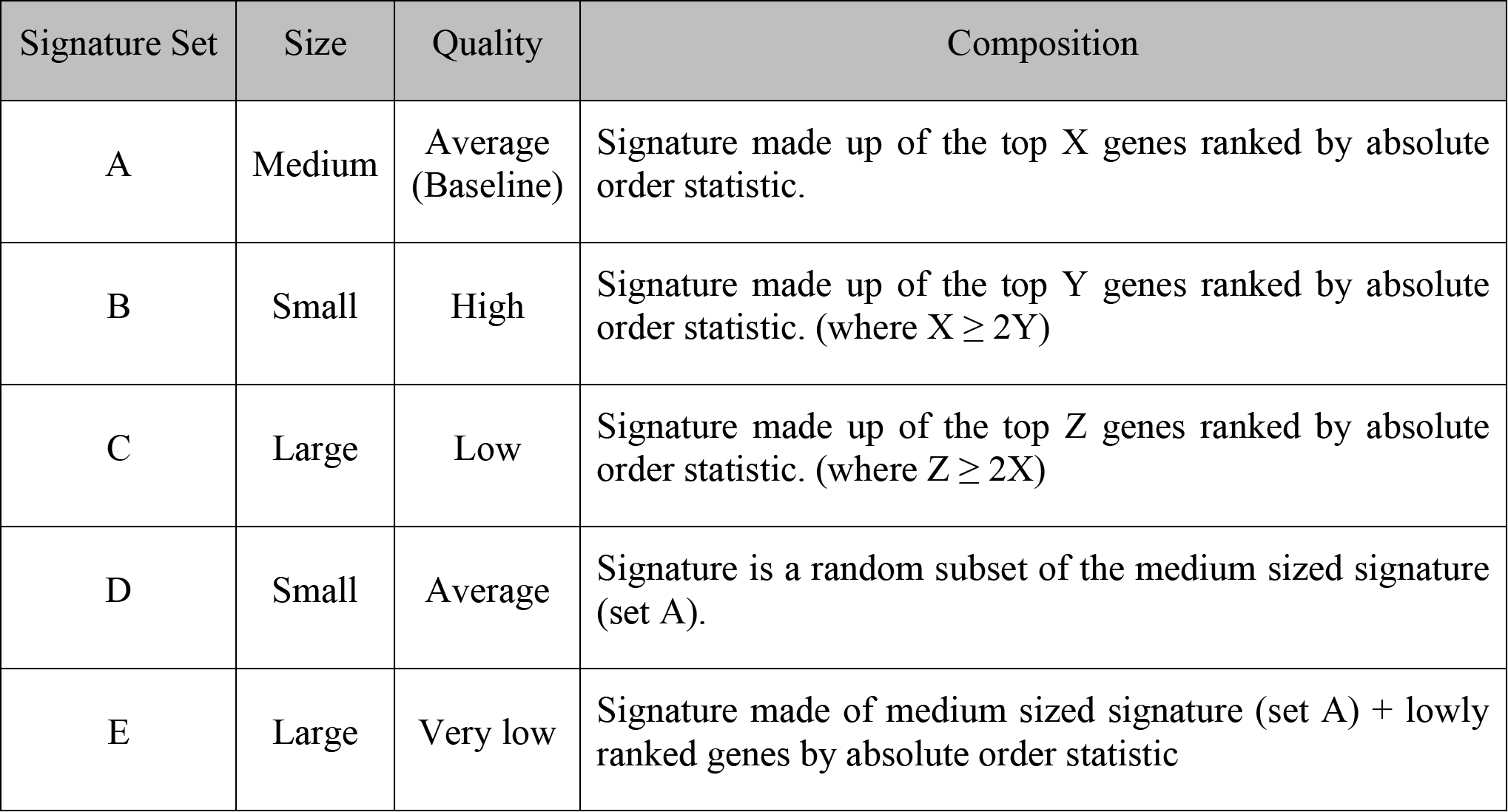
Different quality levels of disease signatures and its composition. Order statistic of a gene: *abs*(*LFC*) × −*log*_10_(*adjusted p* − *value*). X = 250 for gastric cancer analysis; X = 120 for epilepsy analysis.

| Signature Set | Size | Quality | Composition |
| --- | --- | --- | --- |
| A | Medium | Average (Baseline) | Signature made up of the top X genes ranked by absolute order statistic. |
| B | Small | High | Signature made up of the top Y genes ranked by absolute order statistic. (where $X \geq 2Y$) |
| C | Large | Low | Signature made up of the top Z genes ranked by absolute order statistic. (where $Z \geq 2X$) |
| D | Small | Average | Signature is a random subset of the medium sized signature (set A). |
| E | Large | Very low | Signature made of medium sized signature (set A) + lowly ranked genes by absolute order statistic |

For the analysis of set D, the signature of set A was randomly subsetted by one-third 1000 times and two-thirds, also 1000 times, and the drug-disease scores of these signatures were computed. The median and standard deviation of these scores were reported for the analysis of set D. For the analysis of set E, an increasing number of noisy non-DEGs were added to set A signature to dilute the quality of the query signature. We diluted the query signature across four levels, where the number of noisy non-DEGs added is 20%, 50%, 100% and 150% of the size of set A signature. At each dilution level, 1000 different sets of genes were randomly selected from the pool of noisy non-DEGs, and appended to set A. For the gastric cancer study, the noisy non-DEGs are lowly ranked genes which have order statistic < 0.02 (ranked 20,000^th^ and beyond), whereas for the epilepsy study, they are identified by their small absolute LFC and large p-adjusted values (absolute LFC < 1 and q-value > 0.05, order statistics ranked 20,000^th^ and beyond). The drug-disease score, as well as its significance, for each of these noisy signatures were computed. Finally, the number of noisy signatures from set E that still attained a significant score was determined.

### 2.7 Addition of Simulated Noise to LINCS Data

A predetermined amount of noise was added to 50 randomly selected signatures from LINCS, using a simplified method from an earlier work (Dembélé 2013). Briefly, noise is introduced to the level 3 expression values of the 978 landmark genes, using the following model:

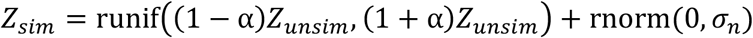

where *Z* is the expression of each gene probe and *α* represents the amount of variance noise added. The LINCS level 3 data was downloaded from the same Clue.io page as above (https://clue.io/data/CMap2020#LINCS2020, (CMap LINCS 2020)), and it comprises over 1.8 million drug instances. The signatures that have been selected for noise simulation are listed in the Supplementary Materials (Table S1).

A user-defined parameter, *λ*, is used to determine the gene average level variation range. This model assumes that the simulated value of each probe is uniformly distributed around the average expression value. The width of the uniform distribution is obtained using the exponential distribution, *α* = *λ*exp (−*λ* × *Z*_unsim_), and is expressed as a percentage of the unsimulated expression value. The use of the exponential distribution to derive the width of gene variation range causes the lowly expressed genes to have larger variability, and vice versa. A second term is added to the simulated data to act as a small additive noise. It is derived from a normal distribution of zero mean and a user-defined standard deviation, *σ*_*n*_. This small additive noise is independent of the gene expression of the probe, and increases with *σ*_*n*_.

Using an identical pipeline as CMap, the simulated expression values of the landmark genes are used to infer the expression of all other genes in LINCS, and finally converted to level 5 data, which represents the replicate-consensus signature in CMap (Subramanian et al. 2017). The amount of eventual noise introduced to a signature is quantified using the spearman rank correlation between the unsimulated and simulated signature.

### 2.8 Comparison by Spearman Correlation of Simulated Signatures

For each signature that has been randomly chosen (termed *randSig*) to be simulated, 1000 (for MCF7, HEK293, NEU and NPC) and 784 (for CD34) signatures from the same cell line were randomly selected, including the unsimulated version of *randSig*, to form the pool of signatures (termed *sigPool*) to be analysed.

First, a query signature was derived from the unsimulated version of *randSig* by subsetting the top 100 genes by absolute LFC. This query signature is a good representation of the transcriptional profile of *randSig*, and was used to calculate similarity scores with the signatures in *sigPool*. The signatures in *sigPool* were then ranked, based on the similarity score, from the strongest phenocopy to the strongest reversal of *randSig* (*sigPoolRank_unsim_*).

Second, noise was introduced to *randSig* to obtain the simulated version of *randSig*. The same filter criteria (i.e. top 100 genes by absolute LFC) was applied to the simulated *randSig*, to generate a simulated query signature to be evaluated against *sigPool*. The signatures in *sigPool* were then ranked based on the new similarity score with the simulated query signature (*sigPoolRank_sim_*).

Finally, the spearman rank correlation between *sigPoolRank_unsim_* and *sigPoolRank_sim_* was determined, for the top and bottom ranked 10% signatures in *sigPoolRank_unsim_*. This analysis was repeated for 50 different versions of simulated *randSig*; and the median correlation was used to compare the performance of the methods at every noise level.

### 2.9 Comparison by AUC of Score Significance of Simulated RandSig

Using the query signature of the unsimulated *randSig*, its similarity score significance with signatures in *sigPool* was also determined. Signatures with significance value smaller than 0.05 were labelled to have a significant relationship with *randSig*, and vice versa; known as ‘actual’ labels.

The score significance of the simulated *randSig* with signatures in *sigPool* is then calculated, to obtain the ‘predicted’ labels. Together, the ‘actual’ and ‘predicted’ labels are used to compute the AUC of the simulated *randSig*. Importantly, if both the ‘actual’ and ‘predicted’ labels are significant, yet the similarity score signs are opposite, the ‘predicted’ label is replaced to be not significant. The median AUC obtained from the 50 simulated *randSig*s were used to evaluate the performance of the methods, at every noise level.

## 3 Results

### 3.1 Comparison by Number of Significantly Scored Signatures

The ability of an algorithm to significantly score a reversal signature (drug-disease) or a true phenocopy signature (drug-drug) is an indicator of its ability to identify true positives. Our first evaluation was based on the effects of 17b-estradiol (E2), which is a natural estrogen receptor ligand. The E2 signature in MCF7 was obtained from an independent study, and it was used as the model signature in the initial CMap publication (Frasor et al. 2004; Lamb et al. 2006). Even though it was reported that there were 40 up and 89 down regulated genes in the E2 signature, we found that only 32 and 67, respectively, of these genes were within LINCS L1000 gene space.

We queried the E2 signature against 357 true phenocopy (ESR agonist) and 273 true reversal (ESR antagonist) drug signatures treated on MCF7 breast cancer cell lines. The E2 signature was also previously used in the benchmarking study by (Lin et al. 2020) and here, we perform the evaluation again using the most recent version of the compound annotations and CMap data. The Zhang method was able to significantly score the highest number of true phenocopy drugs, while the KS and Zhang had similar performance to significantly score true reversal drug signatures of E2. Conversely, the XSum method significantly scored the least number of true phenocopy and true reversal signatures of E2 (Figure 1A).

**Figure 1:**
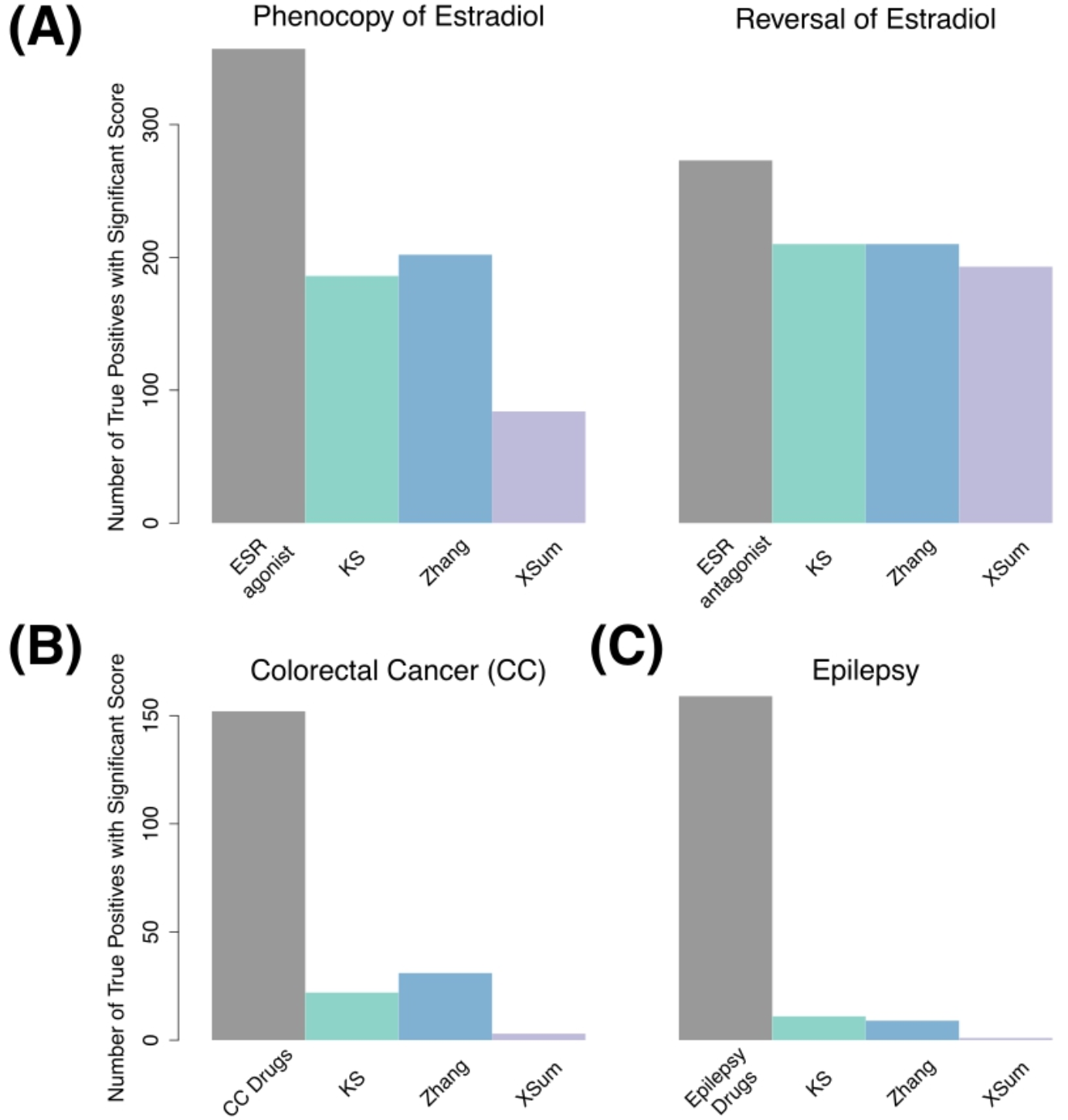
Number of significantly scored signatures for true phenocopy and reversal of 17b-estradiol (A), reversal signatures for colorectal cancer (B) and epilepsy (C).

We next queried a colorectal cancer (CC) signature against 152 HT29 (colon cancer cell line) signatures that were treated with drugs administered to CC patients or compounds that were experimentally validated to reverse CC pathology (Table 3). This CC signature was derived from three independent pairs of comparison between early-stage and metastatic stage colon tumours, and only genes which showed consistent expression changes across the three analyses were compiled to form the CC signature. The compiled CC signature consists of 73 up and 54 down regulated genes (Noort et al. 2014). However, only 60 and 30, respectively, of these genes are found in the LINCS gene space, and queried against the HT29 signatures. The Zhang method had the greatest number of significantly scored signatures for CC, followed by the KS method. The XSum method had the poorest performance, identifying only four out of 152 signatures that used CC reversal drugs (Figure 1B).

For the third analysis, we obtained an epilepsy signature from a pilocarpine-induced status epilepticus (SE) mouse model (Hansen et al. 2014). Post six weeks SE, the RNA-seq evidence suggests increased cellular excitability and morphogenesis in the mice. This epilepsy signature at six weeks was made up of 40 up and 77 down regulated genes (absolute fold change > 2 and q-value < 0.05) in the LINCS gene space. This signature was queried against epilepsy reversal drug signatures in LINCS (NPC and MCF7 cell lines). Additionally, the signatures of sitagliptin, a recent experimentally validated drug that reduced seizure scores in a mouse model of pharmacoresistant epilepsy, were also labelled as reversal signatures of epilepsy (Mirza et al. 2017). Generally, the three methods performed poorly, significantly scoring only less than 7% of all the reversal signatures. Among them, the KS and Zhang methods were able to significantly score a comparable number of reversal signatures, whereas the XSum method had the lowest number of significantly scored reversal signatures (Figure 1C).

Based on the analysis of the E2, colorectal cancer and epilepsy signatures, the Zhang method significantly scored the greatest number of true phenocopy signatures, whereas the KS and Zhang methods significantly scored a comparable number of reversal signatures. On the other hand, the XSum method had the least number of significantly scored true phenocopy and reversal signatures.

### 3.2 Comparison by AUC of Signature Scores

The Receiver Operator Curve (ROC) records how the true positive rate of a predictor model changes with respect to its false positive rate. The method that attains the greatest area-under-the-curve (AUC) suggests that the scores are more indicative of the true drug-disease (or drug-drug) relationship. In the context of drug repurposing, the partial AUC, at lower false positive rate (FPR) thresholds, is also computed (J. Cheng et al. 2014; Lin et al. 2020). The reason is that drug repurposing typically investigates only the top few prioritised drugs, hence a method with higher specificity at low FPR is more desirable.

For the analysis of 17b-estradiol, we queried the E2 signature against 31,471 MCF7 signatures that were treated with 2,325 unique compounds with known targets and MOAs from LINCS. Among these, there are 14 and seven compounds whose MOA was annotated as an ESR agonist and antagonist respectively. The signatures of the 14 ESR agonists are marked as true phenocopy signatures; while the signatures of the 7 ESR antagonists are marked as true reversal signatures. The Zhang method outperforms the other two methods when identifying true phenocopy signatures (AUC – 0.87, AUC0.1 – 0.047, AUC0.01 – 1.8 e-3). For identifying true reversal signatures, the KS and Zhang methods display similar performance and accuracy (KS: AUC – 0.89, AUC0.1 – 0.066, AUC0.01 – 4.2 e-3, Zhang: AUC – 0.90, AUC0.1 – 0.065, AUC0.01 – 4.1 e-3, Table 5). In both analysis for true phenocopy and true reversal signatures of E2, the XSum method attains the lowest AUC.

**Table 5:** AUC of each method in the analysis of 17b-estradiol, colorectal cancer and epilepsy signatures. The bolded values indicate the method which attained the highest AUC.

|  |  | ESR agonists | ESR antagonists | Colorectal Cancer Drugs | Epilepsy Drugs (NPC) | Epilepsy Drugs (MCF7) |
| --- | --- | --- | --- | --- | --- | --- |
| KS | AUC | 0.80 | 0.89 | 0.60 | 0.55 | 0.51 |
|  | AUC0.1 | 0.042 | **0.066** | 9.3 e-3 | 5.1 e-3 | **7.3 e-3** |
|  | AUC0.01 | 1.5 e-3 | **4.2 e-3** | 1.1 e-4 | **1.8 e-4** | **2.2 e-4** |
| Zhang | AUC | **0.87** | **0.90** | **0.65** | 0.52 | **0.53** |
|  | AUC0.1 | **0.047** | 0.065 | **9.9 e-3** | **8.4 e-3** | 5.6 e-3 |
|  | AUC0.01 | **1.8 e-3** | 4.1 e-3 | **1.9 e-4** | 0.9 e-4 | 0.8 e-4 |
| XSum | AUC | 0.81 | 0.80 | 0.57 | **0.57** | 0.47 |
|  | AUC0.1 | 0.028 | 0.030 | 4.2 e-3 | 1.3 e-3 | 1.2 e-3 |
|  | AUC0.01 | 0.5 e-3 | 0.7 e-3 | 0.4 e-4 | 0 | 0 |

Next, we queried the CC signature against 20,866 signatures that were obtained from treating 2,221 unique small drug compounds on HT29 cell lines. Signatures that were generated from CC treatment drugs (Table 3) are marked as reversal signatures. Additionally, the signatures of citalopram and troglitazone, both drugs were reported to significantly reduce tumour volume in mouse model CC, were also marked as reversal signatures (Noort et al. 2014). The Zhang method attained the largest AUC across all FPR thresholds (AUC – 0.64, AUC0.1 – 0.01, AUC0.01 – 1.9 e-4), indicating that the Zhang method is more accurate than the KS and XSum methods when predicting drug-disease relationship in CC (Table 5). The XSum method attained a significantly lower AUC at all false positive rate (FPR) thresholds, suggesting its poor early retrieval performance for the CC signature.

We independently queried the epilepsy signature against 5,638 NPCs and 31,471 MCF7 signatures. Signatures that were perturbed with epilepsy treatment drugs (Table 3), as well as sitagliptin, were marked as reversal signatures of epilepsy. There were 67 and 92 signatures of these compounds in the NPC and MCF7 cell lines respectively. The KS method attained the largest AUC0.01 in both the NPC and MCF7 cell lines (AUC0.01 – 1.8 e-4 (NPC), AUC0.01 – 2.2 e-4 (MCF7)). The Zhang method attained the largest AUC0.1 (AUC0.1 – 8.4 e-3) for signatures in NPC, while the KS method attained the largest AUC0.1 (AUC0.1 – 7.3 e-3) for signatures in MCF7. For the full AUC, the XSum (AUC – 0.57) attained the largest area for NPC signatures, while the Zhang (AUC – 0.53) attained the largest area for MCF7 signatures (Table 5).

Generally, the Zhang method was able to attain the largest AUC when predicting true phenocopy signatures of E2, as well as reversal signatures of CC. The XSum method generally attained the lowest AUC when predicting true phenocopy or reversal signatures. This suggests that the scores from Zhang method, is the best among the three discussed methods, to reflect the true drug-disease (or drug-drug) relationship, whereas the XSum method is the least accurate.

### 3.3 Comparison by Varying Signature Quality

To understand how the drug-disease scores change with respect to the quality of the query signature, we varied the quality of the gastric cancer and epilepsy signatures across five different levels (A–E), and computed the drug-disease score at each quality level. The quality of these signatures were varied by applying different order statistics (*abs*(*LFC*) × −*log*_10_(*adjusted p* − *value*)) cut-off.

Here, we briefly describe each of the five quality levels: Set A to Set C are obtained by first ranking the genes in the DEG analysis by their order statistics. Set A, designated as the baseline for comparison in this analysis, is made up of middle-to-top ranked genes, which are highly and moderately regulated in the disease (top 250 and 120, by order statistics, for gastric cancer and epilepsy, respectively) (Figure 2A, Table 4). Set B comprises only the top few ranked genes and has a high quality, as it is made up of highly regulated genes that define the disease (top 120 and 60, by order statistics, for gastric cancer and epilepsy, respectively). Set C comprises low-to-top ranked genes, and includes mildly regulated genes in addition to Set A (top 500 and 250, by order statistics, for gastric cancer and epilepsy, respectively). As such, Set C is considered to be of low signature quality. Set D is obtained by randomly subsetting baseline set A, and since its proportion of highly-to-moderately regulated genes is comparable with set A, the quality of set D is similar to that of A. The objective of investigating set D scores is to understand the score trend when the query signature size is reduced while its quality is maintained. Set E is generated by including noisy non-DEGs into set A, to form the query signature. As additional noisy non-DEGs are included, the quality of the query signature becomes diluted. Set E explains the score trend when the signature quality is very low (Figure 2A, Table 4).

**Figure 2:**
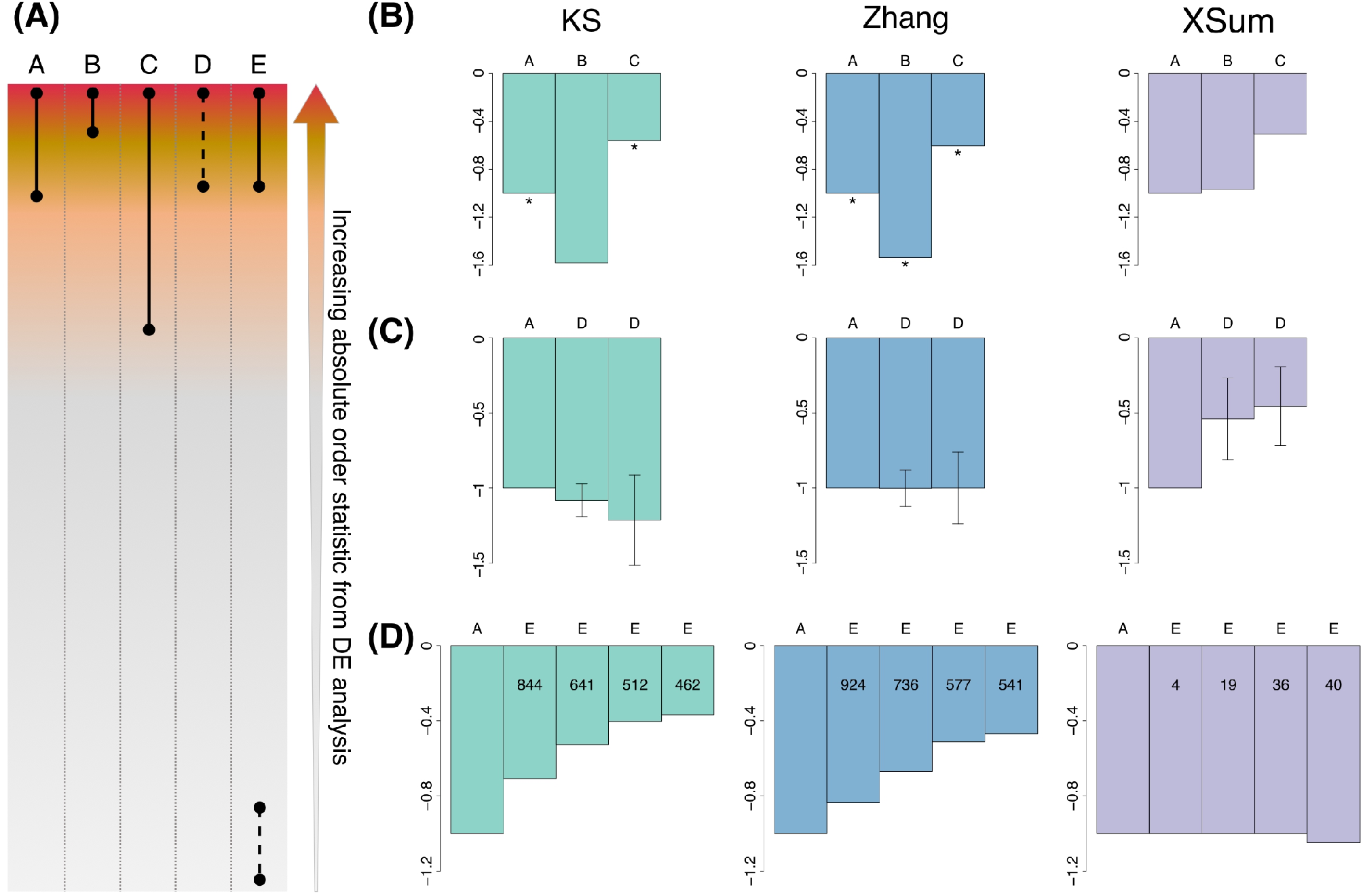
Signature quality in gastric cancer. (A) Schematic illustration of signature composition for Set A to E. Solid lines indicate all genes within the range are included in the query signature, dashed lines indicate a random subset of the genes in the range are included in the query signature (B). Normalised scores by variation of gastric cancer signature quality. Scores are normalised to the baseline set A. Normalised scores for Set A, B and C. Size of set A: 119 genes, set B: 43, set C: 267. * indicates a significant score. (C) Normalised scores for set A and sets D. Size of set A: 119 genes, set D: 78 and 39 respectively. Error bars represent the standard deviation of the scores (n=1000). (D) Normalised scores for set A and sets E. Size of set A: 119 genes, additional number of non-DEGs in set E: 24, 60, 119 and 148 respectively. Internal numerals indicate the number of significant quality-diluted signatures. Green: KS, blue: Zhang, purple: XSum.

Firstly, for the gastric cancer signatures, we queried them against a vorinostat signature in AGS (stomach cancer cell line, 0.12 μM for 24 h — ASG002_AGS_24H:G15). The gastric cancer signature was obtained from the Yonsei database consisting of 65 cancer and 19 normal gastric tissues (Claerhout et al. 2011). An *in vitro* study performed by (Claerhout et al. 2011) indicated positive therapeutic effects of vorinostat towards gastric cancer; therefore we queried its signature against varying signature qualities of gastric cancer. When the quality of gastric cancer signature increases from set A to set B, the drug-disease score becomes more negative for the KS and Zhang methods, indicating that these methods predicted a stronger drug-disease reversal signal. Conversely, as the quality decreases from set A to set C, the KS and Zhang scores become less negative. The Zhang scores obtained across the three sets were also significantly negative, affirming the strong inverse relationship between the vorinostat and gastric cancer signature. The trend of the XSum scores differs from the KS and Zhang methods, as they become less negative across set A, B and C, and the scores are insignificant across the three sets (Figure 2B).

For the analysis of set D for gastric cancer, we generated 1000 random subsets of set A signature. The size of these subset query signatures are approximately two thirds and one third that of set A. The score for each subset signature was calculated, and the median and standard deviation of these 1000 random subset signatures represent the change in drug-disease scores. As the size of the query signature decreases while maintaining the signature quality, only the median of the Zhang scores of set D remain similar to the baseline score of set A. The median KS score of set D is more negative than set A, while the median XSum score is less negative than set A (Figure 2C).

For low quality set E signatures of gastric cancer, we randomly selected noisy non-DEGs and appended them to the baseline set A signature. To investigate how drug-disease scores change as the query signature quality dilutes, we added an increasing number of noisy non-DEGs. The noisy non-DEGs were randomly chosen from the set of genes that are very lowly ranked (absolute order statistics < 0.02) from the GC DEG analysis. For each dilution level, 1000 sets of randomly selected non-DEGs were added to set A signature, and the median drug-disease score obtained from these noisy signatures was determined. Additionally, the score significance for each of the noisy signatures was computed, to determine the number of noisy signatures that attained a significant score.

As the quality of the GC signature dilutes, the median score for KS and Zhang method becomes less negative, while the median XSum score remains generally constant. The KS and Zhang methods have notably more diluted signatures that attained a significant score, than XSum (Figure 2D), suggesting that they can better detect true signals in the query signature in spite of the noisy non-DEGs. The number of noisy signatures that attained a significant score also decreases steadily for the KS and Zhang methods, complying with the reduced performance as the query signature quality decreases.

Next, we varied the quality of the epilepsy signature (Hansen et al. 2014) by applying different order statistics cut-off, in a similar manner as the gastric cancer signature. We queried the aforementioned epilepsy signature against a sitagliptin signature treated in MCF7 cell line (0.125 μM, 24 h). As the quality of the epilepsy signatures increases from set A to set B, the KS and Zhang scores become more negative, and remain significant; while the XSum scores do not change and are insignificant. As the quality decreases from set A to set C, the scores from all three methods become less negative (Figure S1A). Again, these results demonstrate that the KS and Zhang methods produce a stronger drug-disease reversal signal when the query signature has a higher quality.

To obtain set D signatures of epilepsy, we generated 1000 random permutations of two thirds and one thirds of the baseline set A. We computed the score for each signature, as well as their median and standard deviation. As the query signature size decreases while maintaining its quality, only the Zhang score remained stable relative to the baseline score by set A. The median of KS scores becomes more negative, while the median of XSum scores becomes less negative (Figure S1B).

For the analysis of low quality set E signatures of epilepsy, we added an increasing number of noisy non-DEGs of epilepsy to the baseline set A signature of epilepsy, to form the noisy query signature. At each quality dilution level, we generated 1000 such query signatures by randomly selecting noisy non-DEGs (order statistic < 0.1), and computed their drug-disease score. The median KS and Zhang scores become less negative and tend towards zero as more noisy non-DEGs are included, and the number of significant scores also decreases steadily. However, for the XSum method, the median score at each dilution level remained relatively unchanged; and became less negative only when excessively diluted with noisy non-DEGs. Similar to the gastric cancer analysis, the KS and Zhang methods had more significant scores compared to XSum method, suggesting their better capability to detect true signals in the noisy epilepsy query signatures (Figure S1C).

Overall, as the quality of disease signature varies, the KS and Zhang scores change with a similar trend as each other. The changes in the XSum scores generally differ from the other two methods. Table 6 summarises the changes in the scores of all methods, as the quality of the query signature varies.

**Table 6:** Summary of changes in KS, Zhang and XSum scores with respect to disease signatures quality.

| Set (Size) | A (Medium) | B (Small) | C (Large) | D (Small) | E (Large) |
| --- | --- | --- | --- | --- | --- |
| Quality | Baseline | High | Low | Similar to baseline | Very low |
| Trend | Significantly negative? | More negative than (A)? | Less negative than (A) but still significant? | Stable relative to (A)? | Less negative than (A) and no longer significant? |
| KS | Yes | Yes | Yes | No | Yes |
| Zhang | Yes | Yes | Yes | Yes | Yes |
| XSum | No | No | No | No | No |

### 3.4 Correlation between Unsimulated and Simulated Signatures

To further evaluate the robustness of each method to noise, we added noise to the gene expression values of 50 randomly selected level 5 signatures from the LINCS database (*randSig*), across four categorical levels (Figure 3A). These noise levels — termed low, medium, high and extreme — are tuned via two user-defined parameters, *λ* and *σ*_*n*_ (Figure 3B). The *λ* parameter controls the variation range of a gene; whereas *σ*_*n*_ controls the amount of random noise added to each gene. We first introduced noise to the LINCS level 3 instances, and subsequently converted them to their level 5 signatures. The LINCS level 3 data represents the normalised gene expression values for the 978 landmark genes in the L1000 assay for an individual instance, while the level 5 data represents the replicate-consensus signature obtained by a linear combination of the level 3 replicated instances. For each *randSig*, 50 simulated noisy *randSig* were generated, at each noise level.

**Figure 3:**
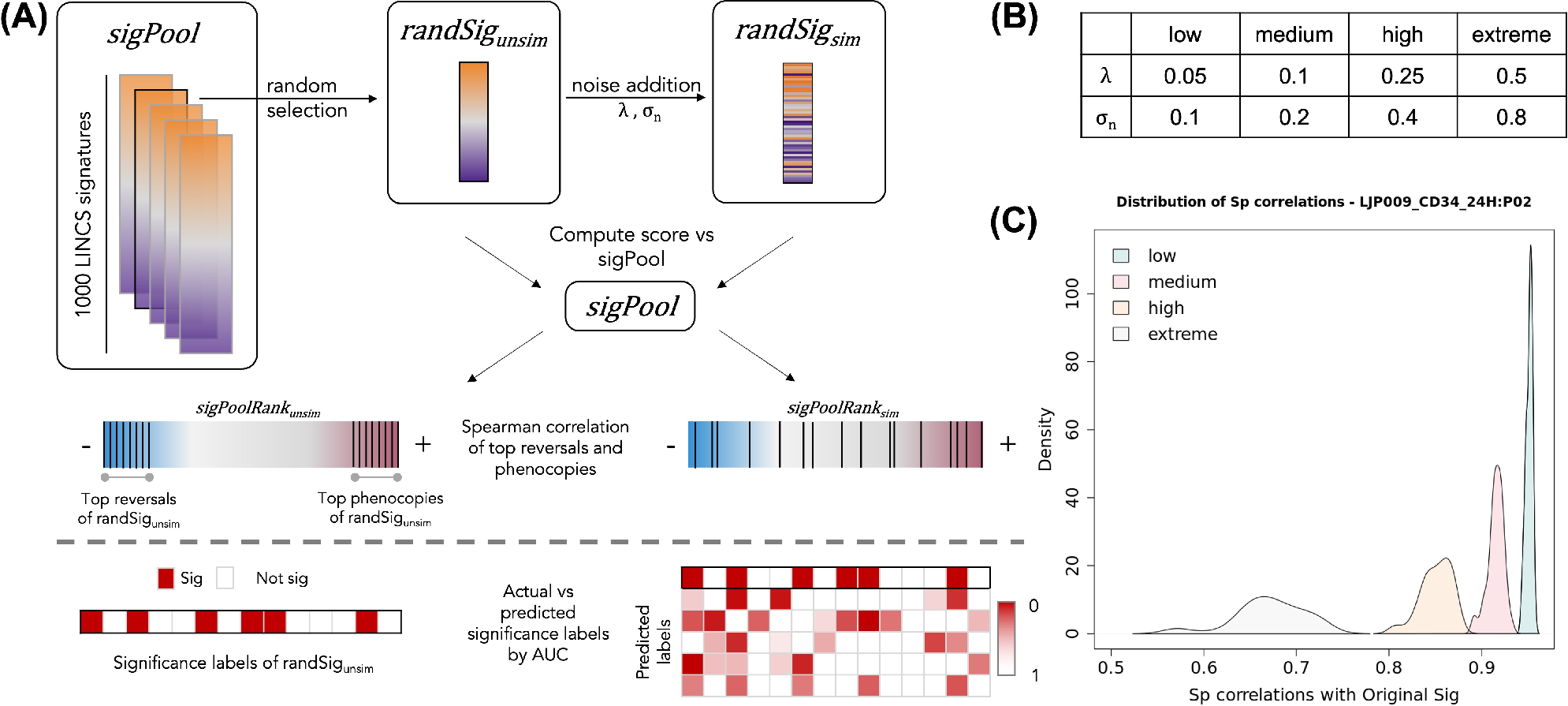
Simulation of noise in differential expression. (A) Schematic of noise addition and analyses of the noisy signatures. (B) Parameters used for noise simulation. (C) Representative density plots of spearman correlations between simulated and unsimulated *randSig* at each noise level (n=50).

The eventual amount of noise that has been added to each *randSig* can be estimated by the spearman rank correlation of the gene expressions, between the unsimulated and simulated *randSig*. We found that the correlation between the unsimulated and low-noise simulated *randSig* is generally above 0.90. As noise level increases, the correlations decrease while the spread of the correlations become wider (Figure 3C). Together, this establishes that noise has been realistically introduced to LINCS data; and at low noise level, the expression profile of the simulated *randSig* is similar to that of the unsimulated *randSig*.

### 3.5 Comparison by Spearman Correlation of Simulated Signature Ranks

The ability of a scoring method to consistently prioritise drug signature in spite of noise within the data, gives an indication to its robustness to noise. To determine the extent to which drug prioritisation has been altered after introduction of noise to *randSig*, we randomly selected a pool of signatures (*sigPool*) from the same cell line as *randSig* in LINCS. We computed the similarity scores between the unsimulated *randSig* and the signatures in *sigPool*, and used them to rank the signatures from the best phenocopy to the best reversal (*sigPoolRank_unsim_*). We then applied the same procedure to each of the 50 simulated *randSig*, to determine the signature ranks after the introduction of noise (*sigPoolRank_sim_*, Figure 3A).

The ranks of the top 10% scoring phenocopy and reversal signatures in *sigPoolRank_unsim_* were evaluated against its new ranks in *sigPoolRank_sim_*. Across noise levels, the correlations for both phenocopy and reversal signatures decrease as noise levels increase (Figure S2A,B). This can be expected as a noisier simulated signature can result in a more drastic change in similarity score, and hence the eventual rank of the signatures. The correlations for phenocopy signatures were also generally higher than that of reversal signatures.

Across methods, the XSum has the most instances in which it attained the highest median correlation for phenocopy signatures, followed by the Zhang method (Figure 4A). For reversal signatures, the KS and Zhang methods have a comparable number of instances that attained the highest median correlation (Figure 4B). It is observed that for most *randSig*s, the method that attained the best correlation at low noise level also has the best correlation at higher noise levels. This suggests that the best performing method is highly dependent on the *randSig* of choice.

**Figure 4:**
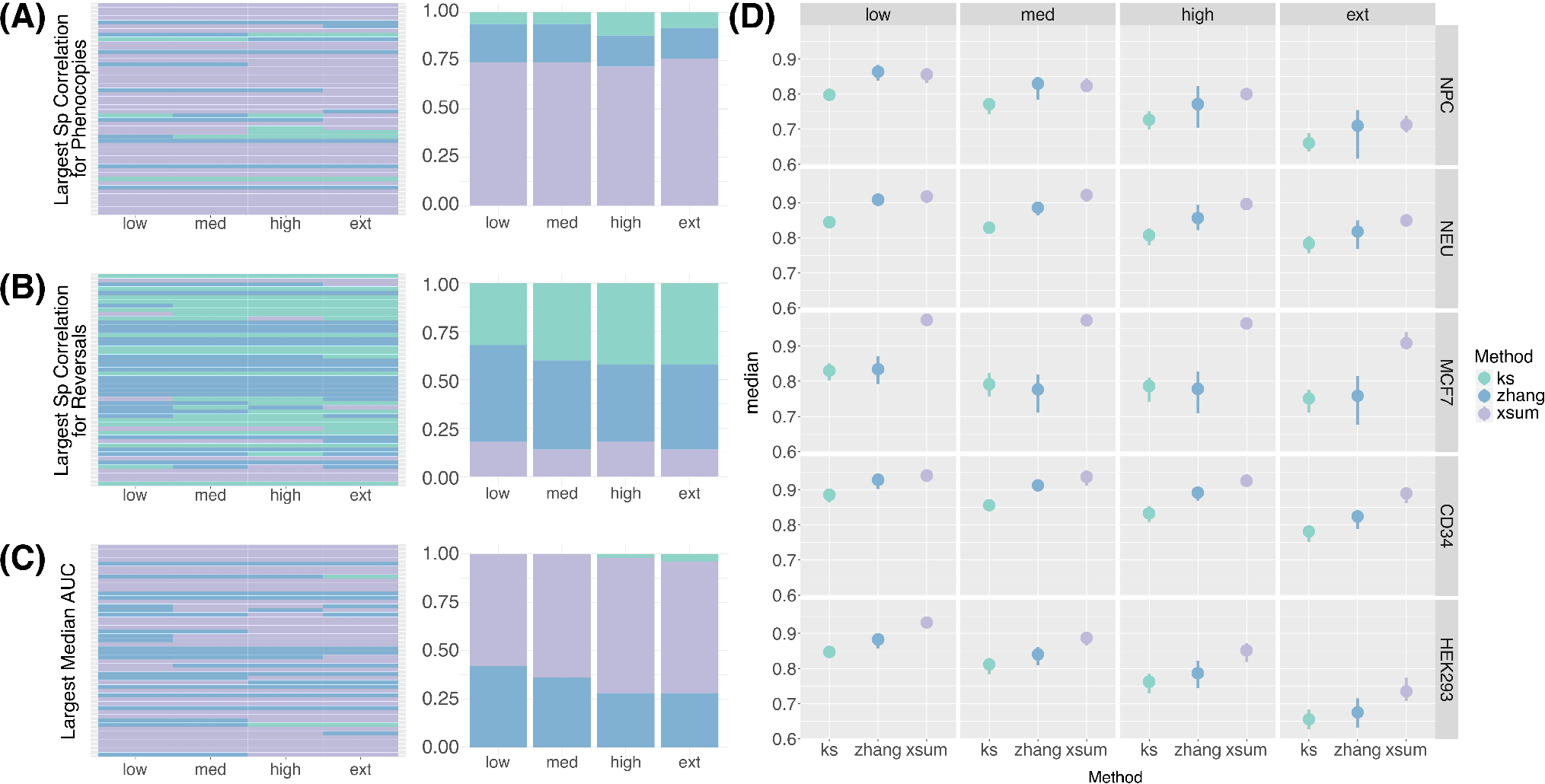
Spearman Correlation of Simulated Signature Ranks. (A) Heatmap (left) and proportion plot (right) indicating the method that attained the largest median correlation for phenocopy signature ranks. The rows of the heatmap represent the *randSig*s that were selected for noise simulation. (B) Heatmap (left) and proportion plot (right) indicating the method that attained the largest median correlation for reversal signature ranks. (C) Heatmap (left) and proportion plot (right) indicating the method that attained the largest median AUC. (D) Point range plot of the median AUC (n=50) between the actual and predicted significance labels of *randSig*. The AUCs of a representative signature from each cell line was used to generate the plot. The complete plot for every *randSig* can be found in the Supplementary Materials. Green: KS, blue: Zhang, purple: XSum.

### 3.6 Comparison by AUC of Score Significance of Simulated RandSig

The second approach we employed to investigate the robustness of the methods, was to determine if the methods retain the significance labels after the inclusion of noise. We first determined the ‘actual’ labels based on the score significance of the unsimulated *randSig*. Next, the score significance of the simulated *randSig*s are computed, to obtain the ‘predicted’ labels. Finally, we compare the ‘actual’ and ‘predicted’ labels to compute the AUC by score significance. A larger AUC indicates that the method is able to significantly score a signature in spite of noise within the data.

Similar to the comparison by spearman correlations, the XSum method has the most instances in which it attained the highest AUC (Figure 4C). The KS method attained the smallest AUC in 41 out of the 50 low-noise simulations, highlighting its inferiority when handling noisy data. As the amount of simulated noise increases from low to extreme, the AUC decreases slightly, indicating that the accuracy of the significance label drops for all three methods (Figure S2C). The relative performance of the methods at low-noise level is also reflective of their relative performance at higher noise levels (Figure 4D, S2C).

## 4 Discussion

In this study, we evaluated the relative performance of three primary in silico drug repurposing methods — KS, Zhang and XSum. The methods are compared based on how sensitive they are to true positives, how their scores change with varying signature quality, and how robust they are to noisy DEG data. The analysis was based on gastric and colorectal cancer signatures, as well as epilepsy signatures in the mouse model, queried against the most recent release of LINCS data. Besides disease signatures, the methods were also compared on their ability to quantify drug-drug similarity using the transcriptional signature of 17b-estradiol.

First, in comparing the methods based on their sensitivity, the XSum method generally has an inferior performance to the KS and Zhang methods. XSum method has the lowest number of significantly scored phenocopy or reversal signatures, as well as the smallest AUC by signature score across the three methods. Between KS and Zhang methods, the latter performed marginally better, notably in identifying phenocopy signatures of E2 and colorectal cancer treatment drug signatures. Our findings from the analysis of E2 signatures with the most recent version of LINCS data supports the results of an earlier study (Lin et al. 2020).

Next, we incorporated noise into the query signature by varying its DEG composition. This gives an insight into how the drug-disease scores change with respect to the quality of query signatures. Although frequently acknowledged to be of great relevance, the influence of the query signature quality on drug-disease indication has been understudied (Zhang and Gant 2008; J. Cheng et al. 2014; Musa et al. 2018). On the basis that a higher quality query signature will produce a stronger drug-disease indication (and vice versa), the results show that the Zhang method best follows this expected change in drug-disease score. The quality of the query signature is approximated to the proportion of highly DEG, based on the order statistic of the genes in a DEG analysis. As the quality of the query signature increases, the magnitude of both KS and Zhang scores increase, suggesting both methods are able to predict a stronger drug-disease reversal potential. When the size of the query signature was reduced while retaining its quality level, only the Zhang scores remained unchanged. The query signature quality was then further diluted by including noisy non-DEGs; the KS and Zhang scores gradually decreased in magnitude and tend towards zero. These methods also detected true signals (true DEGs), within query signatures that have been noised with non-DEGs, better than the XSum method.

Lastly, we simulated noise in the query signature arising from the quantification of gene expressions. We used the spearman rank correlation to evaluate if the scoring methods were able to replicate the drug prioritisation ranks, before and after noise addition in the query signature. For the prioritisation of phenocopy signatures, XSum had a superior performance over the other two methods, and attained the highest median correlation in nearly 75% of the randomised signatures (*randSig*s). For the prioritisation of reversal signatures, there is no one method that consistently outperformed the other methods — all methods have a comparable number of *randSig*s in which it attained the best median correlation. We observed in the analysis of spearman correlation for the top phenocopy and reversal, generally, the method that has the best performance at low noise, is also the best performing method at higher noise levels (78% and 72% of *randSig*s, for top phenocopy and reversal, respectively). This suggests that the best performing method is highly dependent on the *randSig* that has been chosen for simulation.

We further evaluated the ability of the methods to retain significant/non-significant labels after noise addition using the AUC metric. Among the three methods, XSum had the highest median AUC in 60% of the simulations, while the Zhang method had the highest median AUC in 40% of the simulations. There were no simulated signature in which the KS method had the best median AUC. A likely reasoning for the XSum method performing best in most instances, is that the composition of the extreme genes was perturbed to a lesser extent than the actual ranks of the genes. The results from this AUC analysis also show that the relative performance of the methods at low noise is generally reflective of their relative performance at higher noise levels. Together, for the analysis of noise simulation in the query signature, the performance of the Zhang and XSum methods outperforms the KS method, for the ranking of phenocopy signatures and AUC comparisons.

Considering all the comparison metrics, the performance of the three methods are summarised in Figure 5. Our findings suggest that the Zhang method may be most suitable, among the three methods, to predict drug-disease indications with LINCS data. Besides displaying good sensitivity to significantly score true positive signatures, our results show that the Zhang method is robust to noisy query signatures when the composition of DEGs are varied. In terms of noise related to the quantification of gene expressions, the XSum method had most instances in which it performed best, but the Zhang method did not trail far behind. There may be additional factors in the randomly chosen signatures that affect which method performs best with the addition of noise. While out of the scope of this work here, future studies should investigate the qualities of a signature which may dictate the optimal approach.

**Figure 5:**
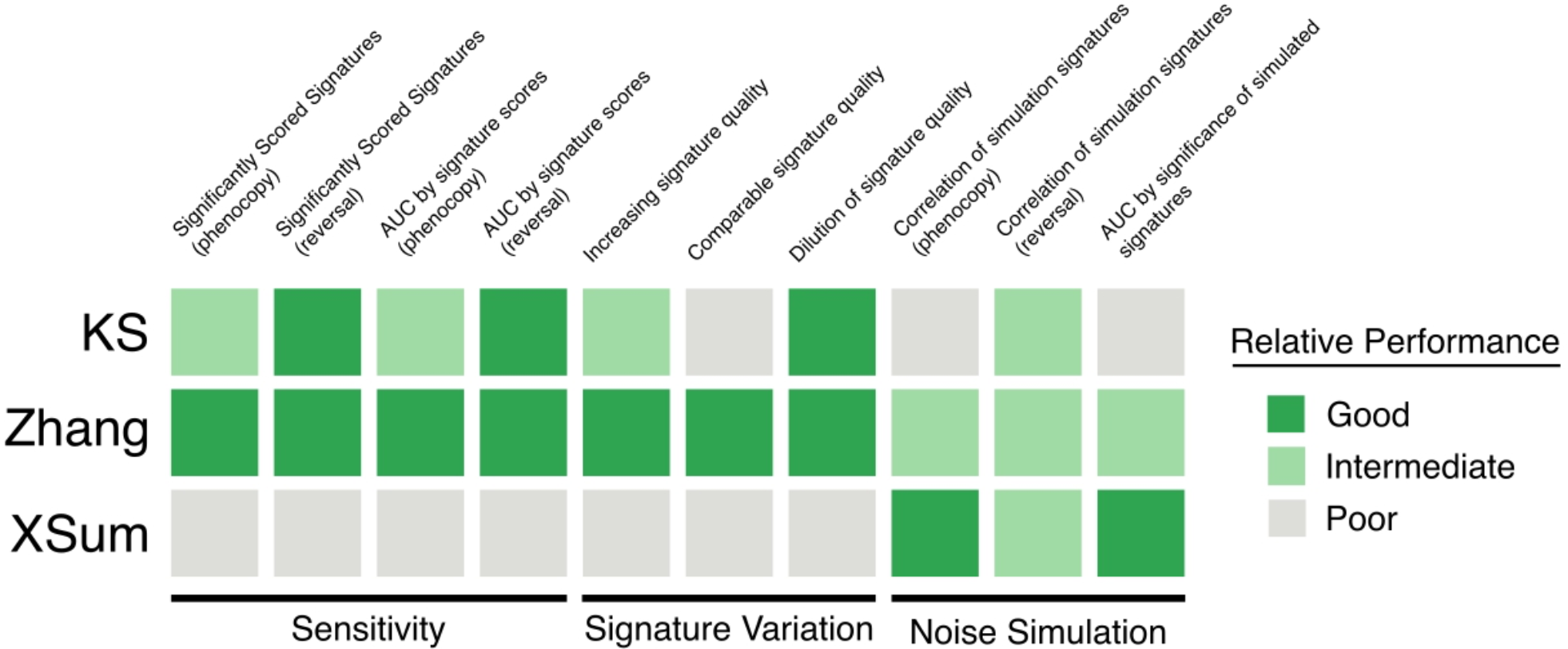
Summary of the relative performance of the three in silico drug repurposing algorithms. Bright green: good, dull green: intermediate, grey: poor.

As noise was introduced to query signatures from the LINCS database in this study, its principles were relevant to the LINCS L1000 array platform. It will be ideal to perform a similar noise simulation study to query signatures obtained using RNA-seq platforms, to understand if the findings of this study extend to noisy RNA-seq derived signatures. Further, this study encompasses two different disease types (cancer and epilepsy) and their related cell lines; it remains to be explored how the performance of the methods may be altered for in silico drug repurposing in other disease types.

This study presents an assessment of three primary in silico drug repurposing methods — KS, Zhang and XSum. These methods are evaluated based on their sensitivity, and performance as query signature quality changes. The Zhang method displays superior sensitivity, especially for phenocopy signatures. For reversal signatures, the KS and Zhang methods have comparable sensitivity. When the quality of query signature is altered by filtering for only top DEGs or by including non-DEGs, the Zhang scores performed as expected with respect to the query size and strength. When the quality of query signature is altered by noising gene expression, the XSum method, followed by the Zhang, best retains information from the original randomly selected signature. The results also suggest that the method which is most robust to noisy gene expression data, may be dependent on additional factors in the original signature. For the signatures used in this study, the XSum and Zhang methods seem to fare better than the KS method.

Together, this work provides guidelines to understand the suitability of three connectivity methods for in silico drug repurposing. It proposes a novel framework to study how connectivity scores are affected by query signature quality, an aspect of in silico drug repurposing that is widely recognised yet understudied.

## 1 Data Availability Statement

Publicly available datasets were analysed in this study. The LINCS data was retrieved from the CMap LINCS Resource 2020 (URL: https://clue.io/data/CMap2020#LINCS2020). The epilepsy signature was retrieved from the Gene Expression Omnibus (GEO) database repository, accession number GSE72402 (URL: https://www.ncbi.nlm.nih.gov/geo/query/acc.cgi?acc=GSE72402). The gastric cancer signature was retrieved from GEO, accession number GSE13861 (URL: https://www.ncbi.nlm.nih.gov/geo/query/acc.cgi?acc=GSE13861). The processed colorectal cancer signature was retrieved from Supplementary Table S3 of (Noort et al. 2014) (URL: https://doi.org/10.1158/0008-5472.CAN-13-3540). The 17b-estradiol signature was obtained from Supplementary Material 1132939s_sigS2. (URL: https://doi.org/10.1126/science.1132939).

## 2 Author Contributions

NT and SRL conceived and designed the project. NT performed and SRL supervised the analysis. NT and SRL wrote the manuscript and all authors approved the final manuscript.

## 4 Acknowledgments

We would like to acknowledge the members of the Integrative Biology of Disease group at the Lee Kong Chian School of Medicine for their helpful comments and suggestions on this work. The computational work for this article was partially performed on resources of the National Supercomputing Centre, Singapore (https://www.nscc.sg).

