## Supplementary Materials for "Evaluating the Robustness of Connectivity Methods to Noise for In Silico Drug Repurposing Studies"

### Supplementary Material

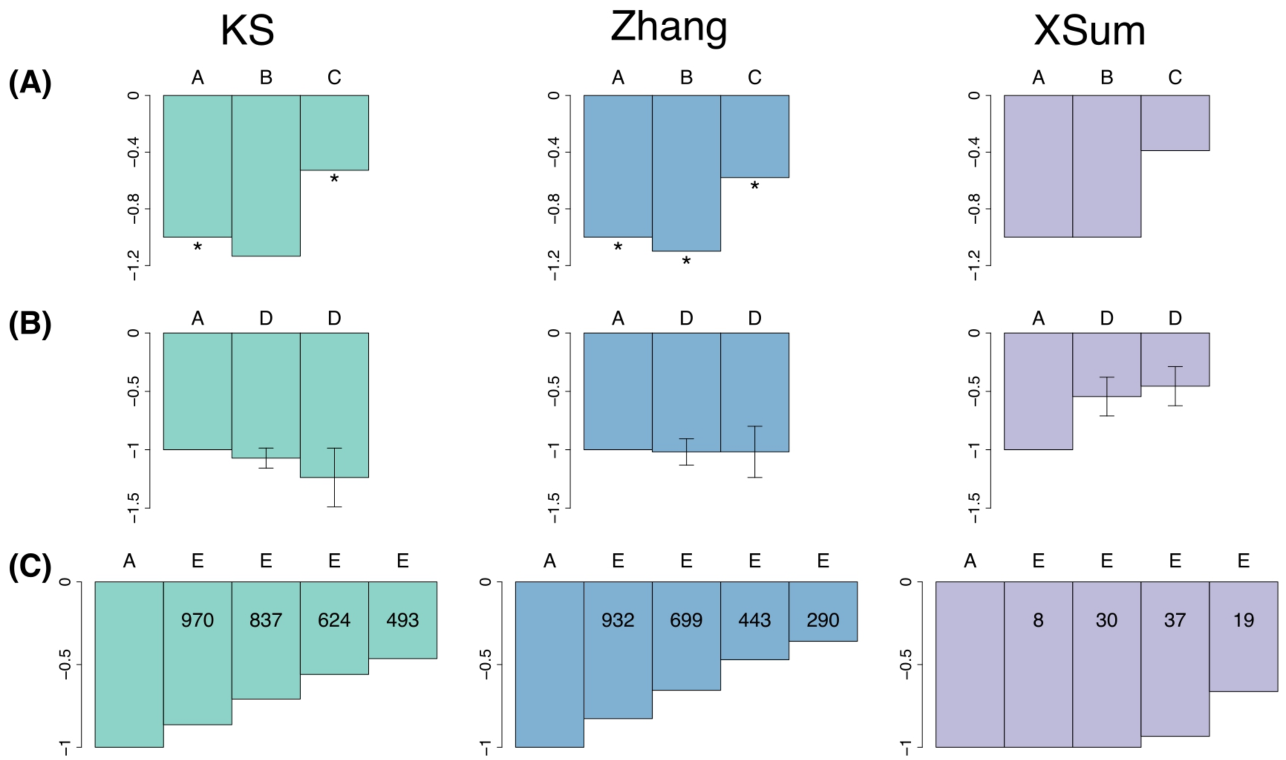


**Supplementary Figure 1**. Effect of signature quality in repurposing for epilepsy. (A) Normalised scores by variation of epilepsy signature quality. Scores are normalised to the baseline set A. Normalised scores for Set A, B and C. Size of set A: 117 genes, set B: 57, set C: 295. * indicates a significant score. (B) Normalised scores for set A and sets D. Size of set A: 117 genes, set D: 78 and 39 respectively. Error bars represent the standard deviation of the scores (n=1000). (C) Normalised scores for set A and sets E. Size of set A: 117 genes, additional number of non-DEGs in set E: 23, 58, 117 and 178 respectively. Internal numerals indicate the number of significant diluted-quality signatures. Green: KS, blue: Zhang, purple: XSum.

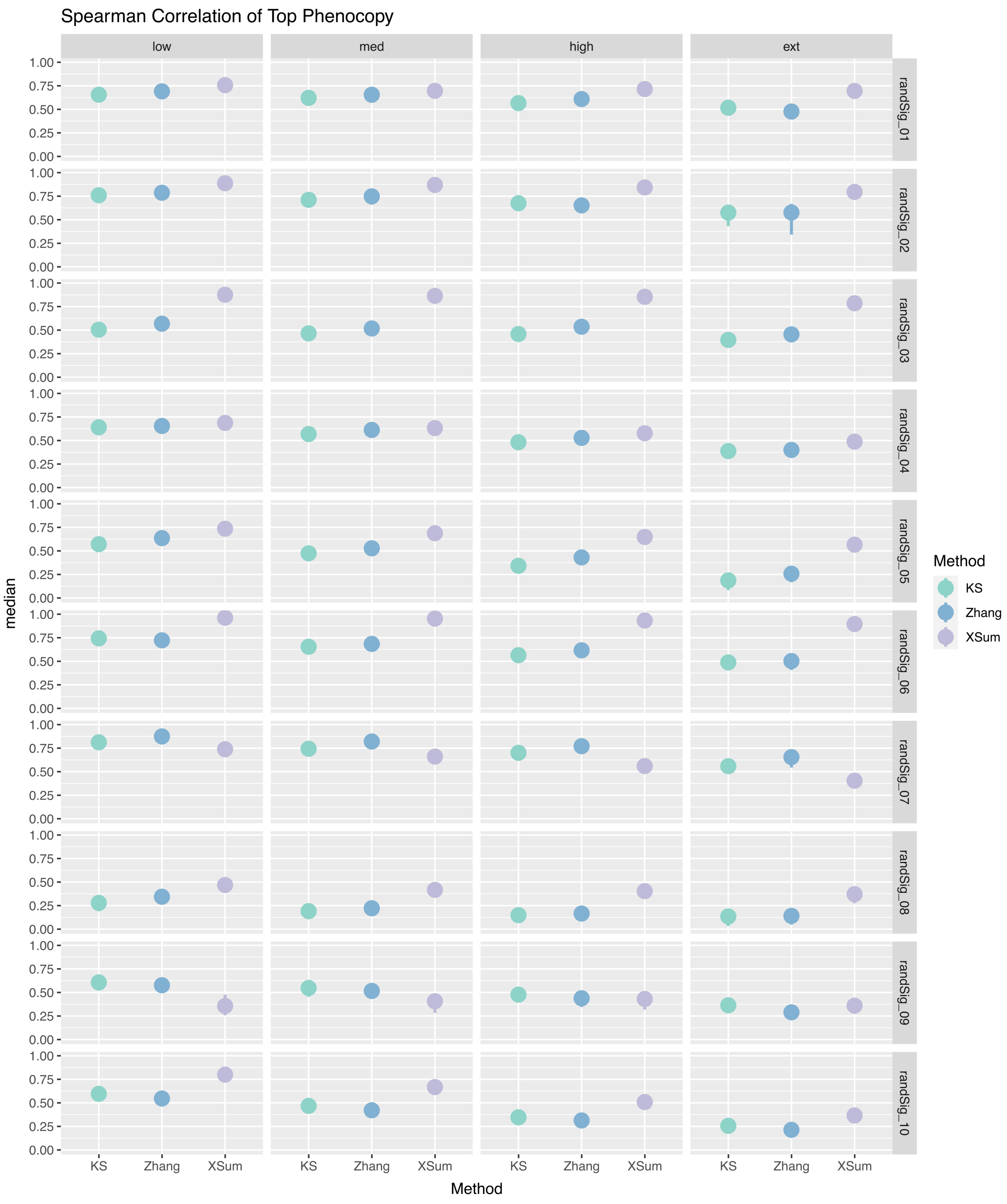
Spearman Correlation of Top Phenocopy
low
med
high
ext
randSig_01
randSig_02
randSig_03
randSig_04
randSig_05
randSig_06
randSig_07
randSig_08
randSig_09
randSig_10
1.00
0.75
0.50
0.25
0.00
median
KS
Zhang
XSum
Method
Method
KS
Zhang
XSum

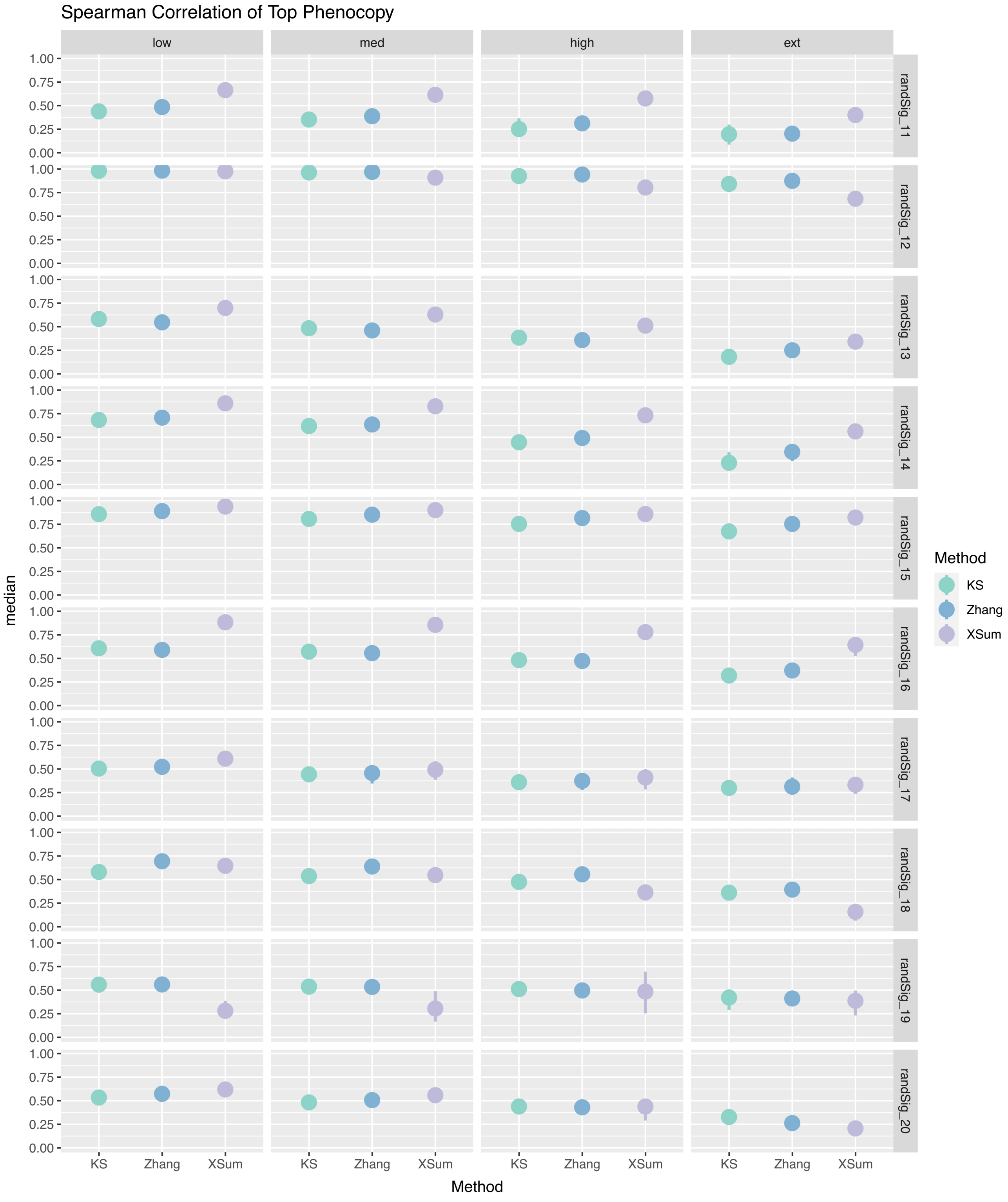
Spearman Correlation of Top Phenocopy
low
med
high
ext
randSig_11
randSig_12
randSig_13
randSig_14
randSig_15
randSig_16
randSig_17
randSig_18
randSig_19
randSig_20
median
1.00
0.75
0.50
0.25
0.00
KS
Zhang
XSum
Method
Method
KS
Zhang
XSum

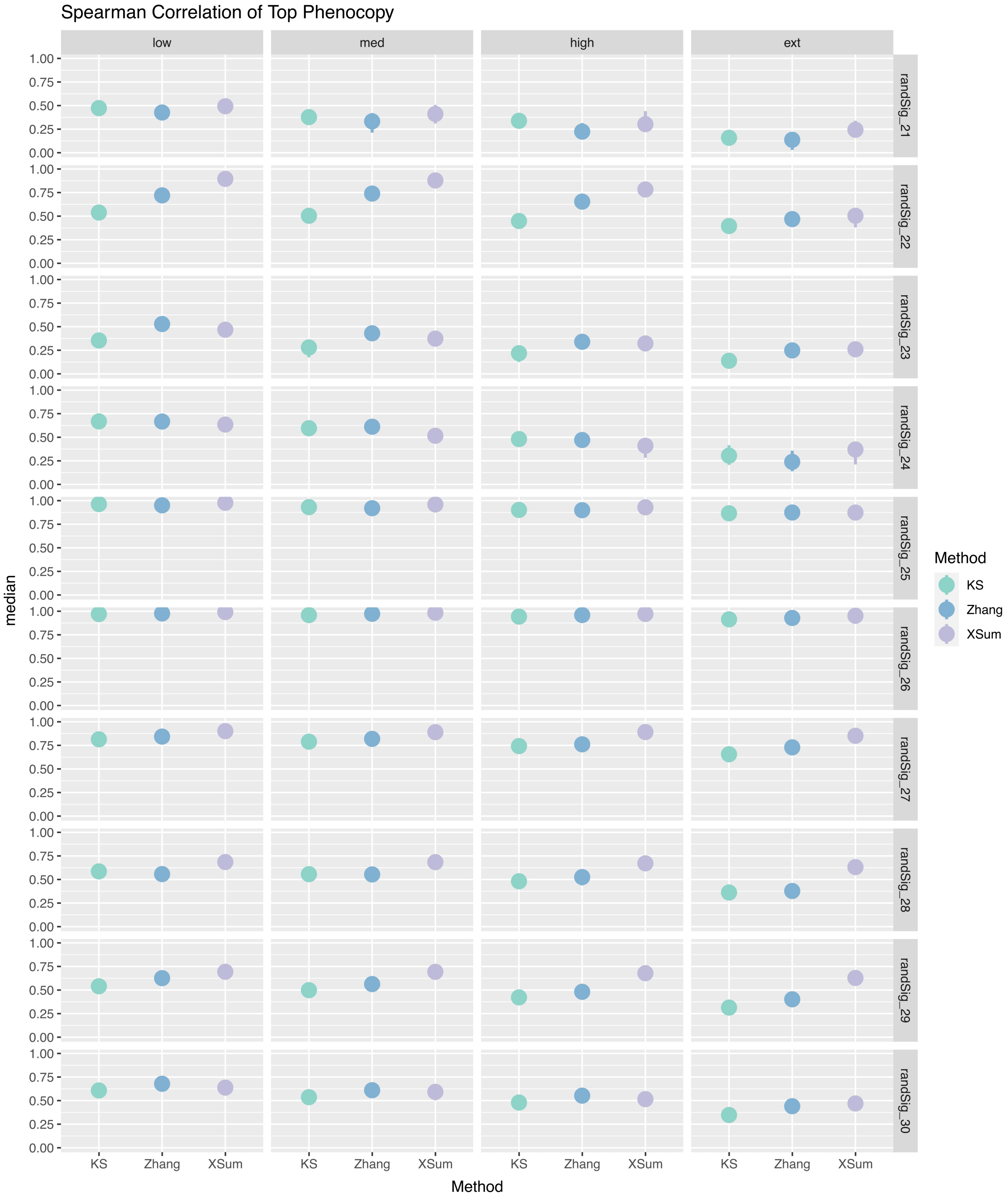

Spearman Correlation of Top Phenocopy
low
med
high
ext
randSig_21
randSig_22
randSig_23
randSig_24
randSig_25
randSig_26
randSig_27
randSig_28
randSig_29
randSig_30
1.00
0.75
0.50
0.25
0.00
median
KS
Zhang
XSum
Method
Method
KS
Zhang
XSum

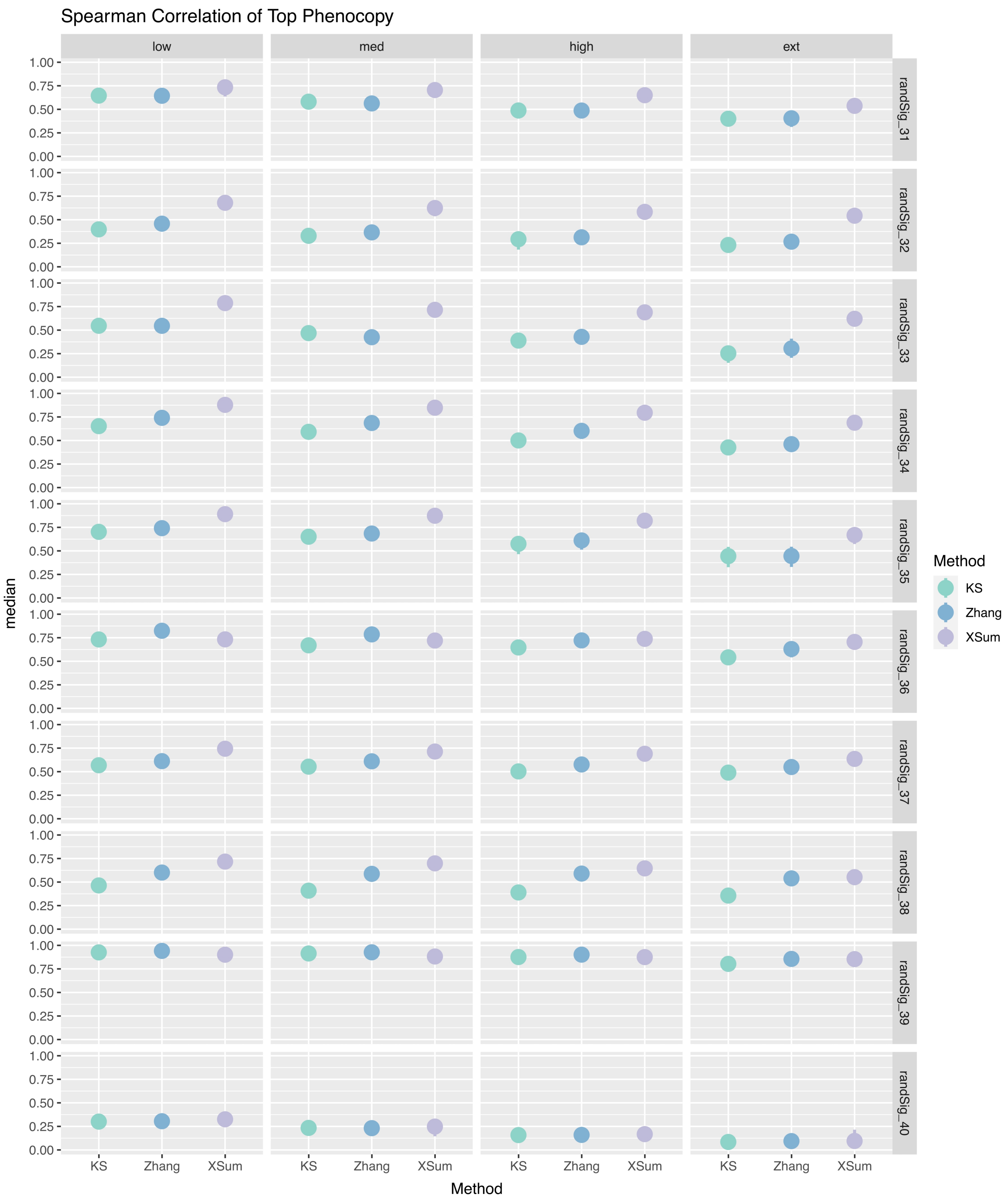

Spearman Correlation of Top Phenocopy
low
med
high
ext
randSig_31
randSig_32
randSig_33
randSig_34
randSig_35
randSig_36
randSig_37
randSig_38
randSig_39
randSig_40
median
1.00
0.75
0.50
0.25
0.00
KS
Zhang
XSum
Method
Method
KS
Zhang
XSum

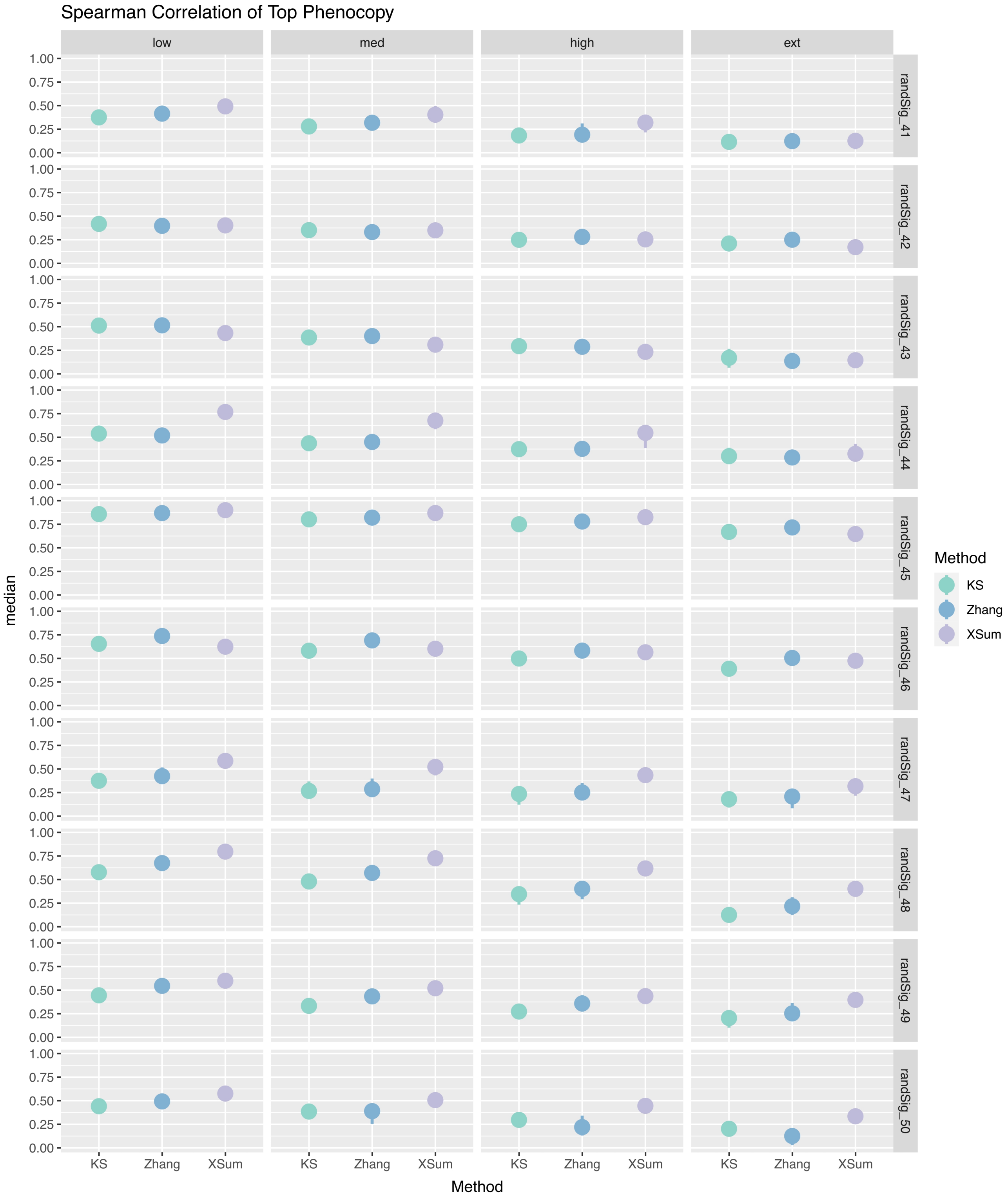


**Supplementary Figure 2A**. Point range plots of the median spearman correlation (n=50) between the unsimulated and simulated ranks for the top phenocopies of randSig.

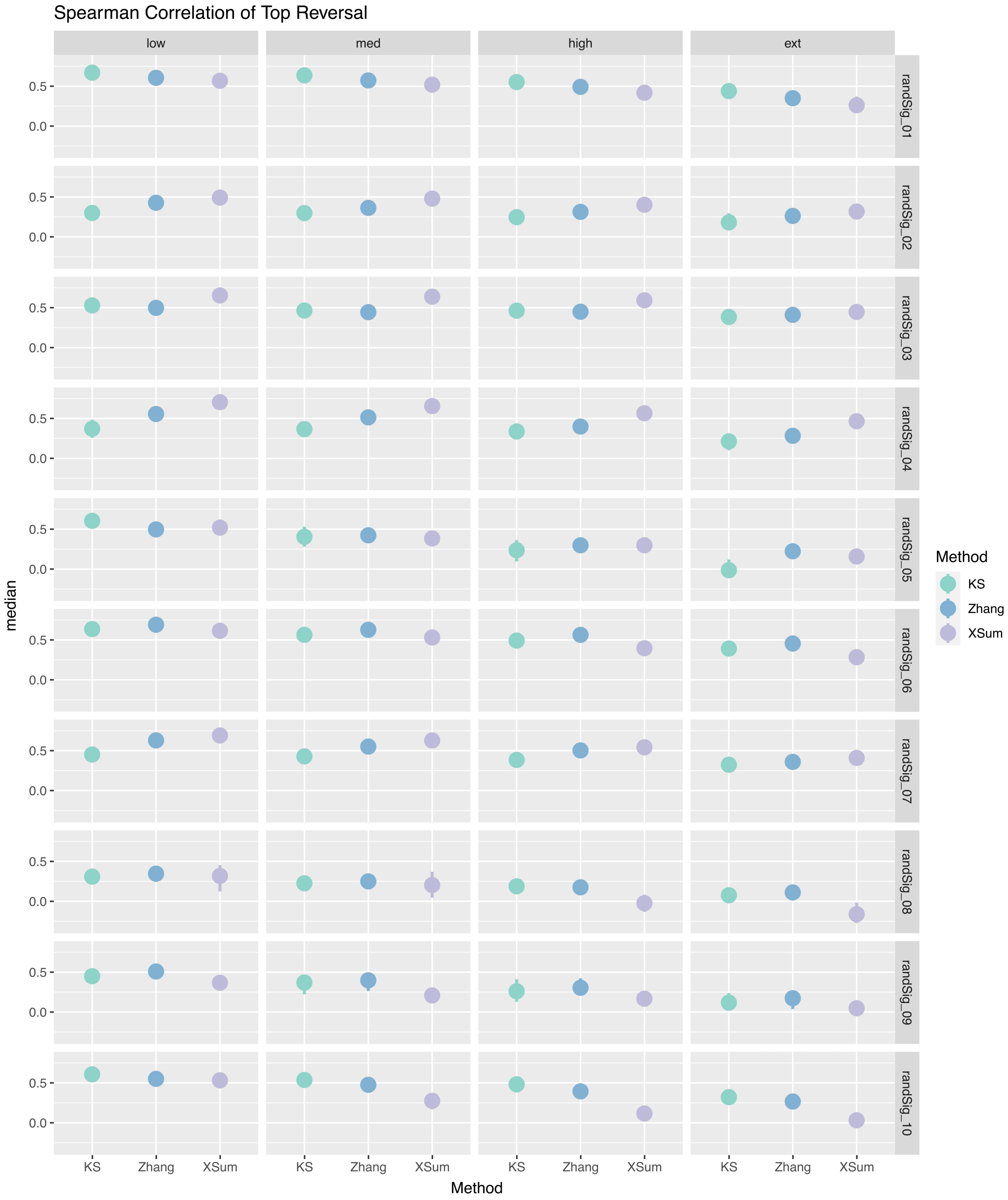

Spearman Correlation of Top Reversal
low
med
high
ext
randSig_01
randSig_02
randSig_03
randSig_04
randSig_05
randSig_06
randSig_07
randSig_08
randSig_09
randSig_10
0.5
0.0
median
KS
Zhang
XSum
Method

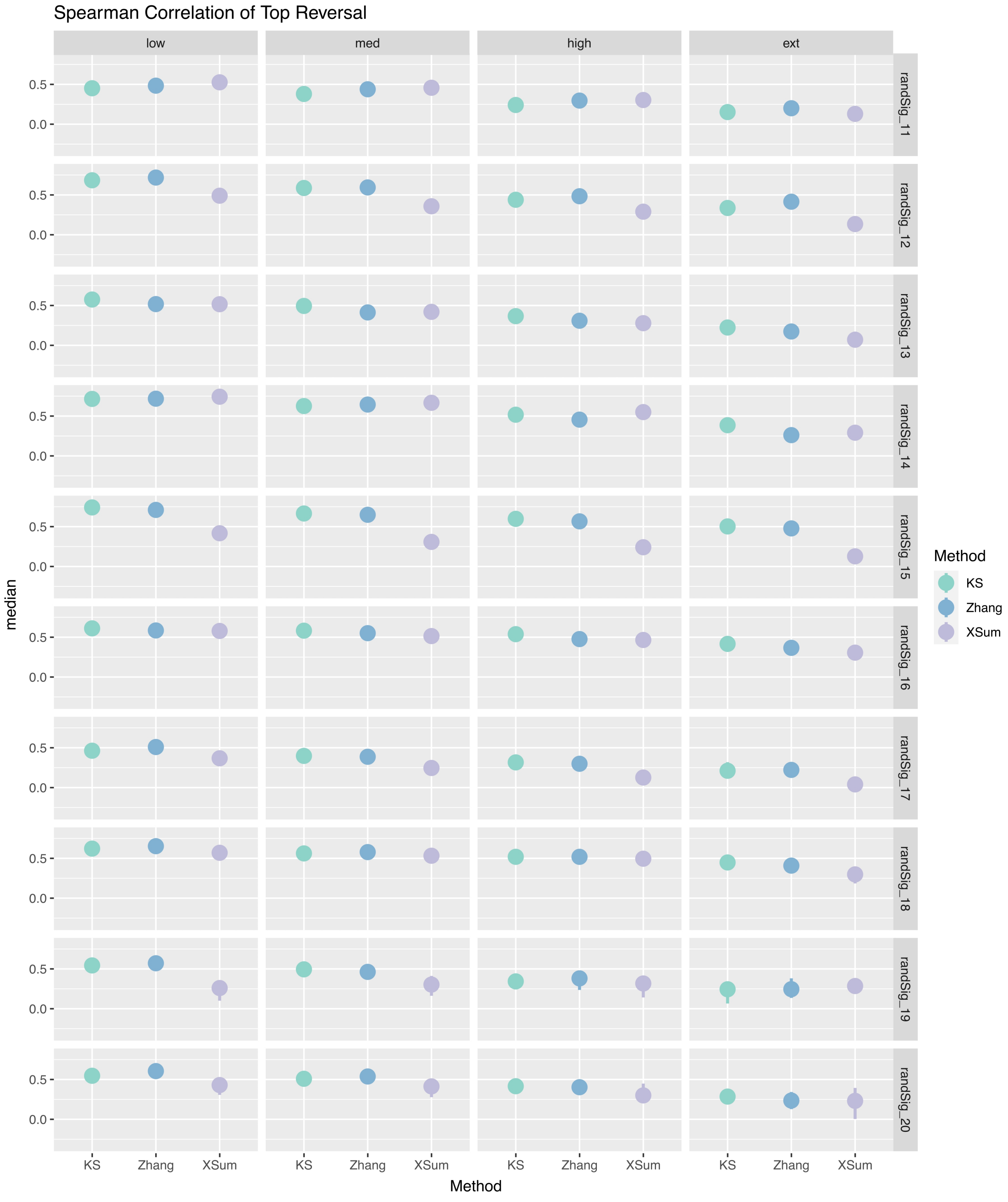
Spearman Correlation of Top Reversal
low
med
high
ext
randSig_11
randSig_12
randSig_13
randSig_14
randSig_15
randSig_16
randSig_17
randSig_18
randSig_19
randSig_20
0.5
0.0
median
KS
Zhang
XSum
Method
Method
KS
Zhang
XSum

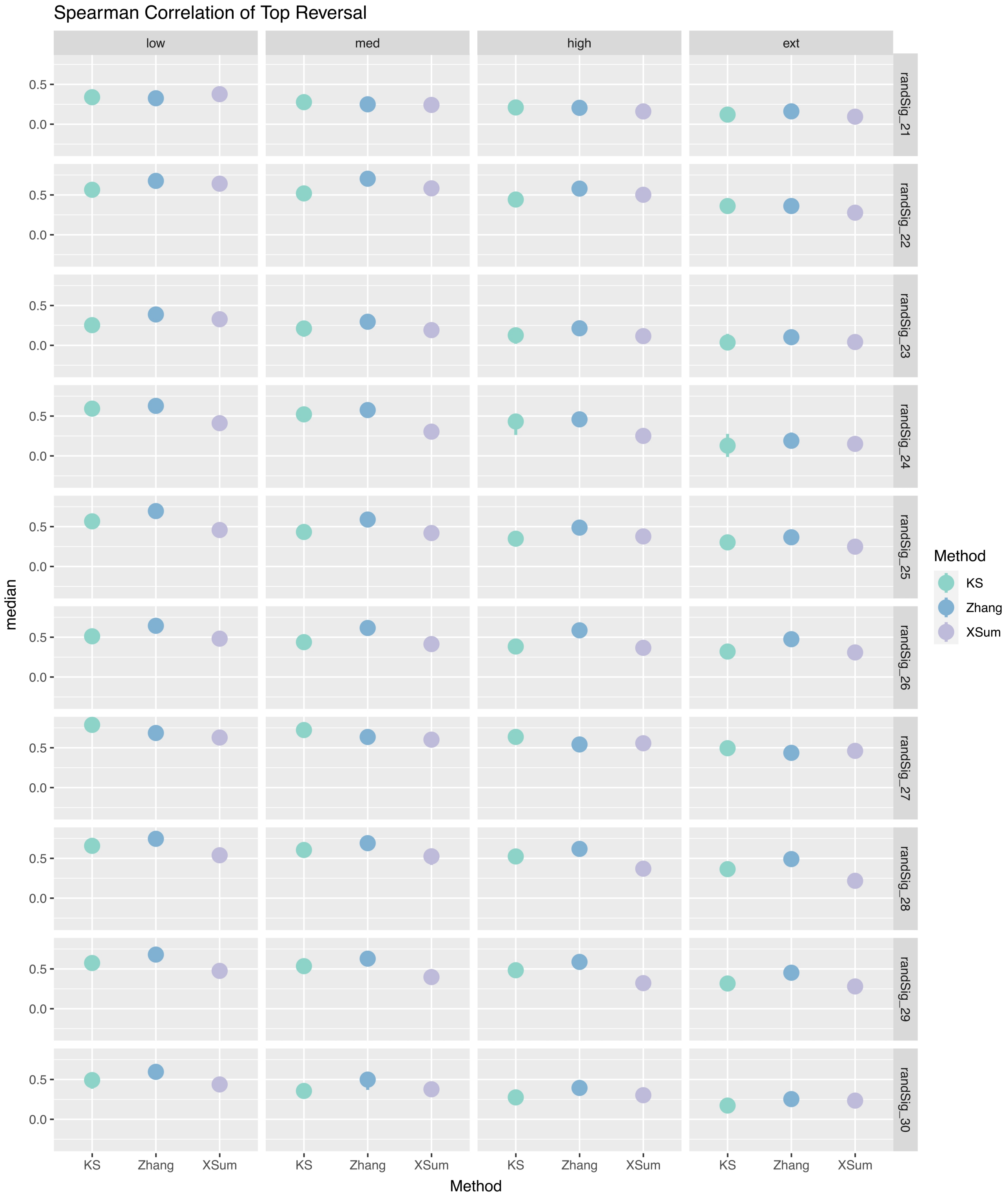
Spearman Correlation of Top Reversal
low
med
high
ext
randSig_21
randSig_22
randSig_23
randSig_24
randSig_25
randSig_26
randSig_27
randSig_28
randSig_29
randSig_30
0.5
0.0
median
KS
Zhang
XSum
Method
Method
KS
Zhang
XSum

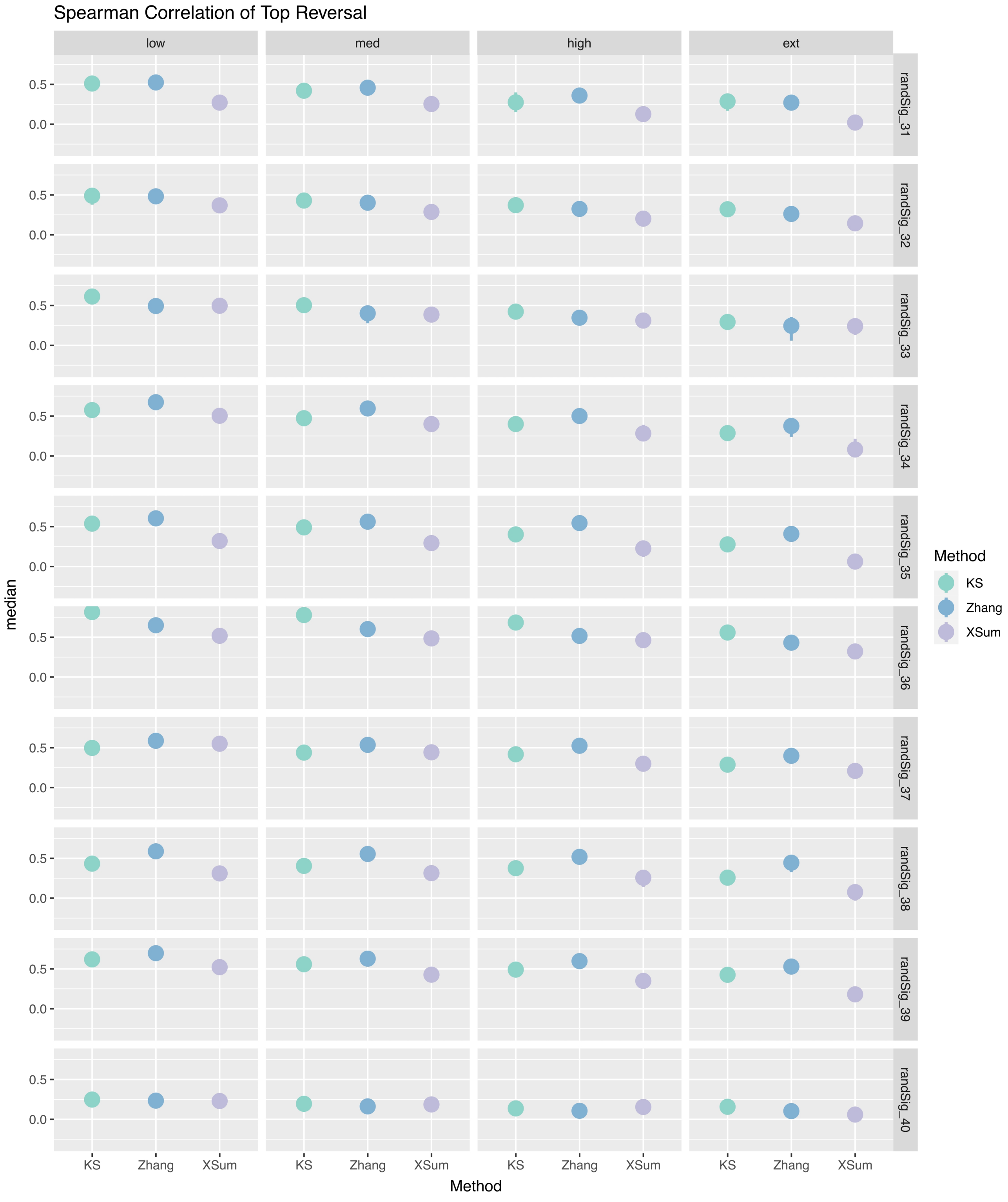
Spearman Correlation of Top Reversal
low
med
high
ext
randSig_31
randSig_32
randSig_33
randSig_34
randSig_35
randSig_36
randSig_37
randSig_38
randSig_39
randSig_40
0.5
0.0
median
KS
Zhang
XSum
Method
Method
KS
Zhang
XSum

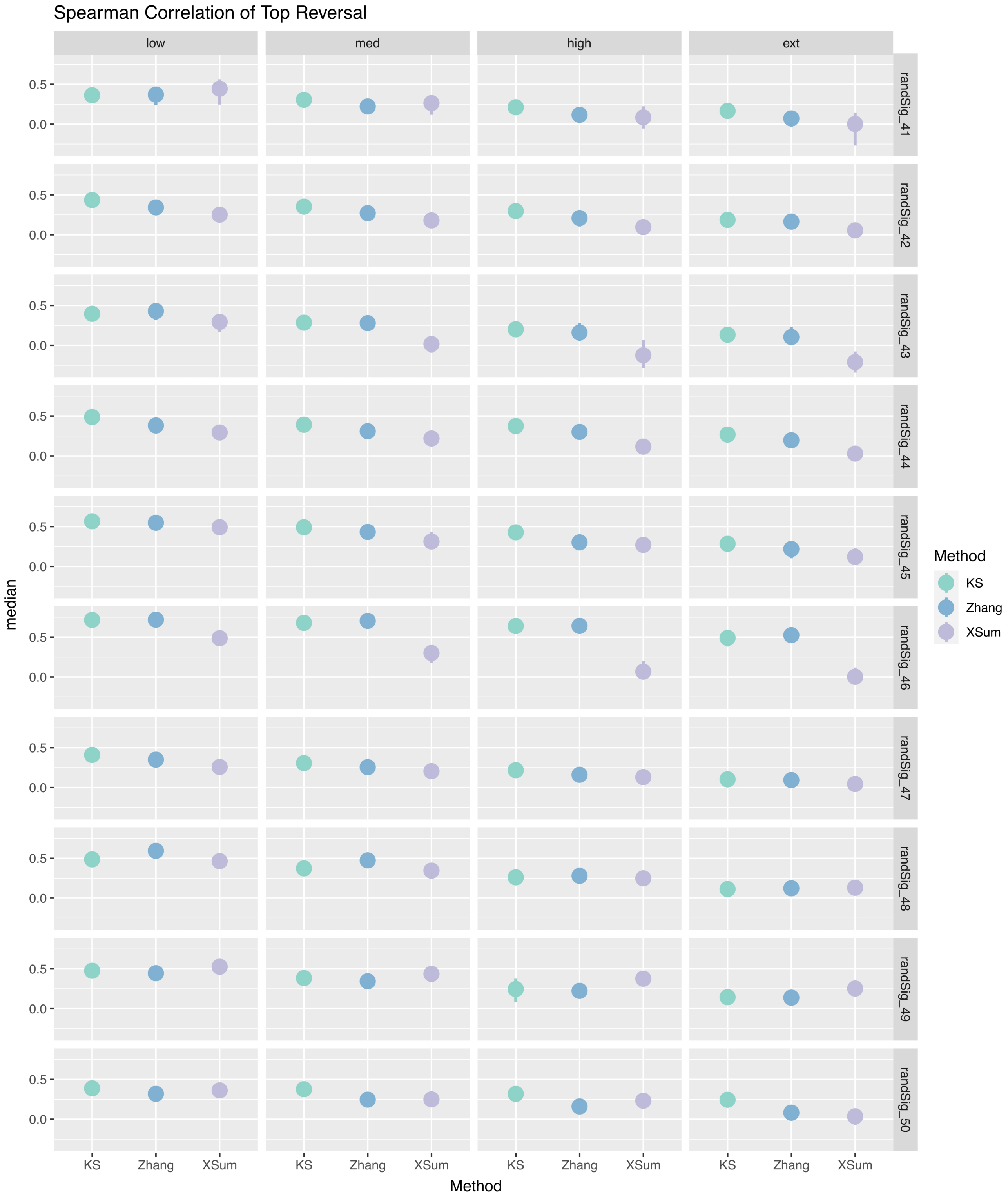


**Supplementary Figure 2B**. Point range plots of the median spearman correlation (n=50) between the unsimulated and simulated ranks for the top reversals of randSig.

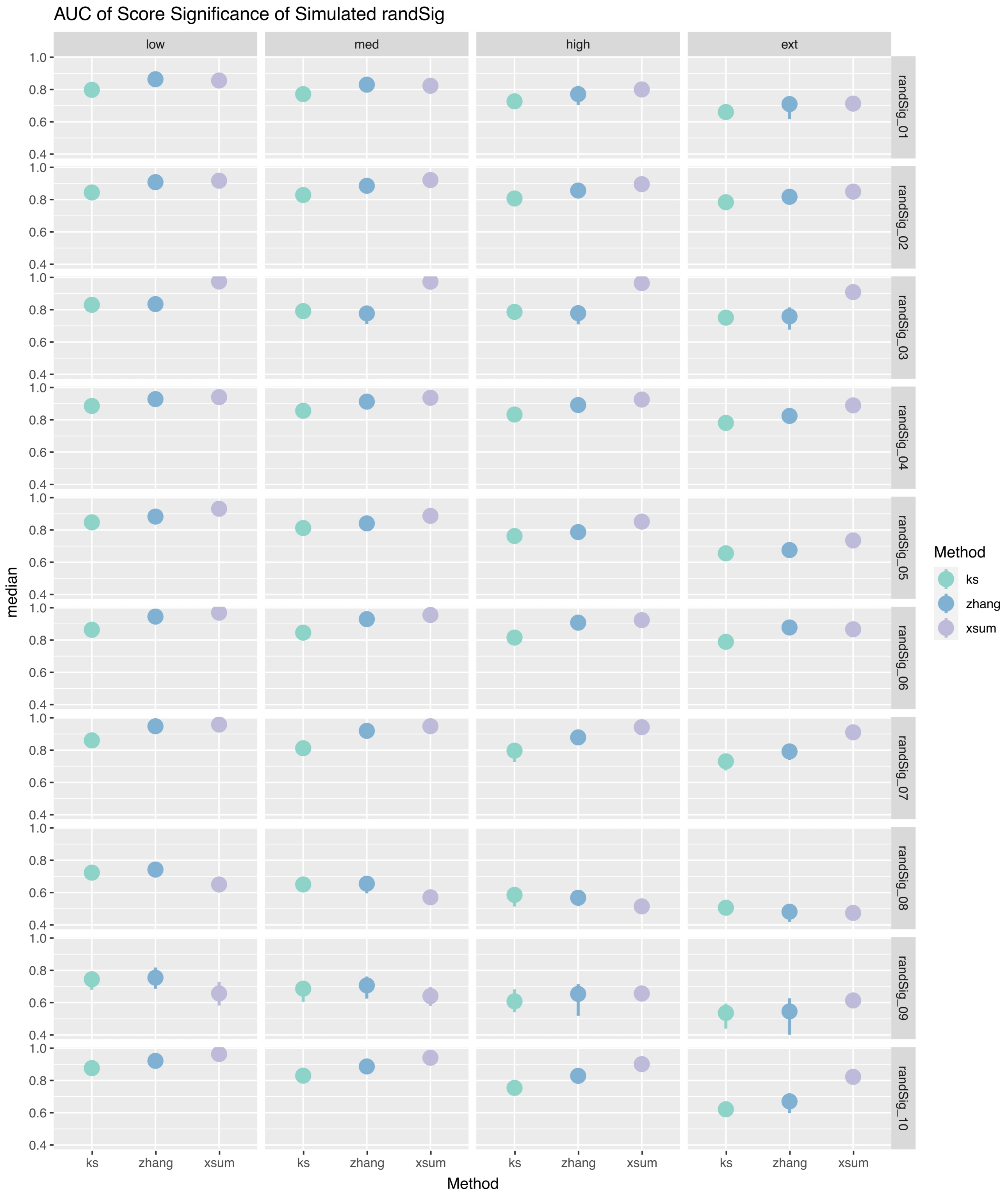
AUC of Score Significance of Simulated randSig
low
med
high
ext
randSig_01
randSig_02
randSig_03
randSig_04
randSig_05
randSig_06
randSig_07
randSig_08
randSig_09
randSig_10
1.0
0.8
0.6
0.4
median
ks
zhang
xsum
Method
Method
ks
zhang
xsum

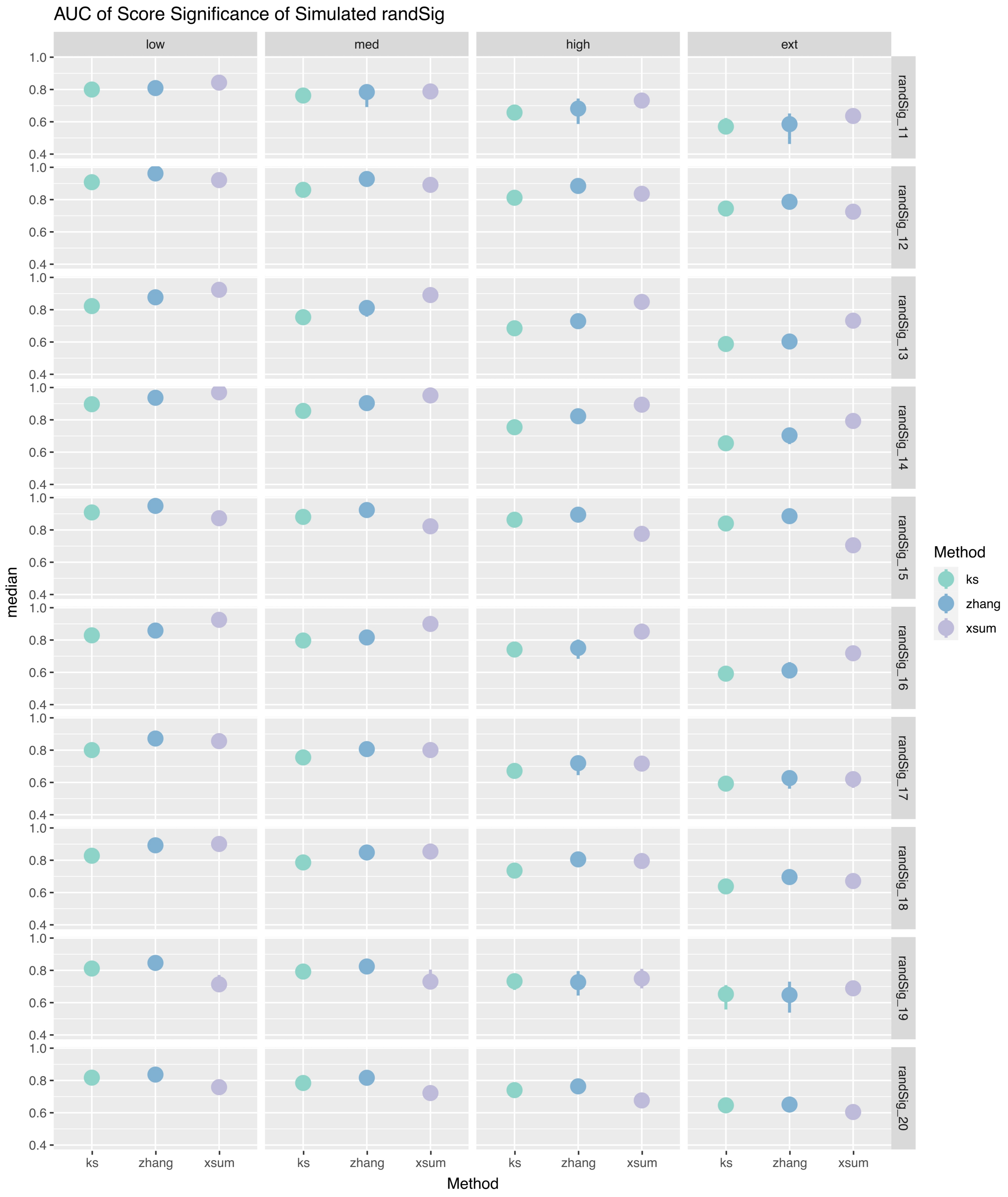
AUC of Score Significance of Simulated randSig
low
med
high
ext
randSig_11
randSig_12
randSig_13
randSig_14
randSig_15
randSig_16
randSig_17
randSig_18
randSig_19
randSig_20
median
1.0
0.8
0.6
0.4
ks
zhang
xsum
Method
Method
ks
zhang
xsum

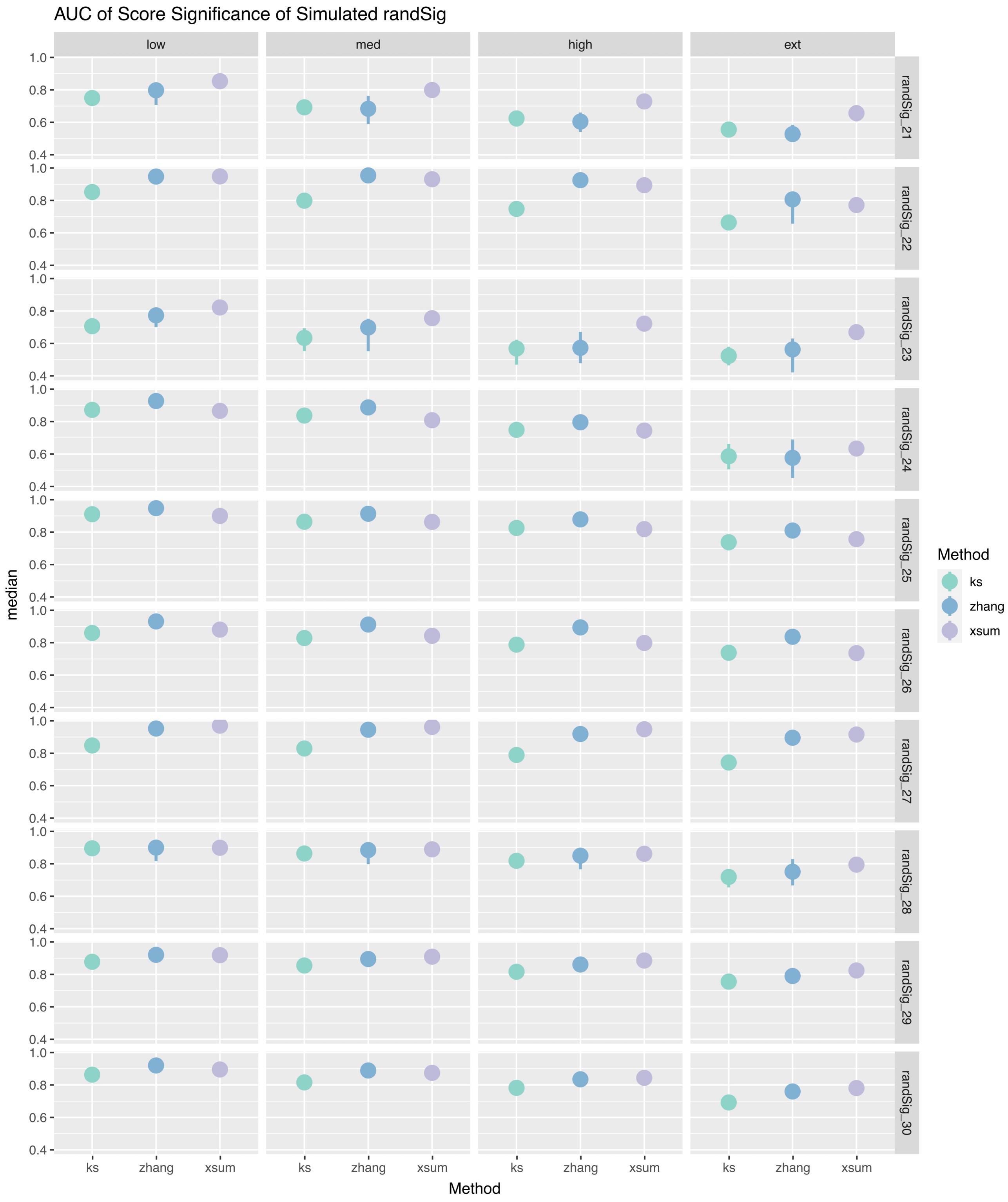
AUC of Score Significance of Simulated randSig
low
med
high
ext
randSig_21
randSig_22
randSig_23
randSig_24
randSig_25
randSig_26
randSig_27
randSig_28
randSig_29
randSig_30
median
1.0
0.8
0.6
0.4
ks
zhang
xsum
Method
Method
ks
zhang
xsum

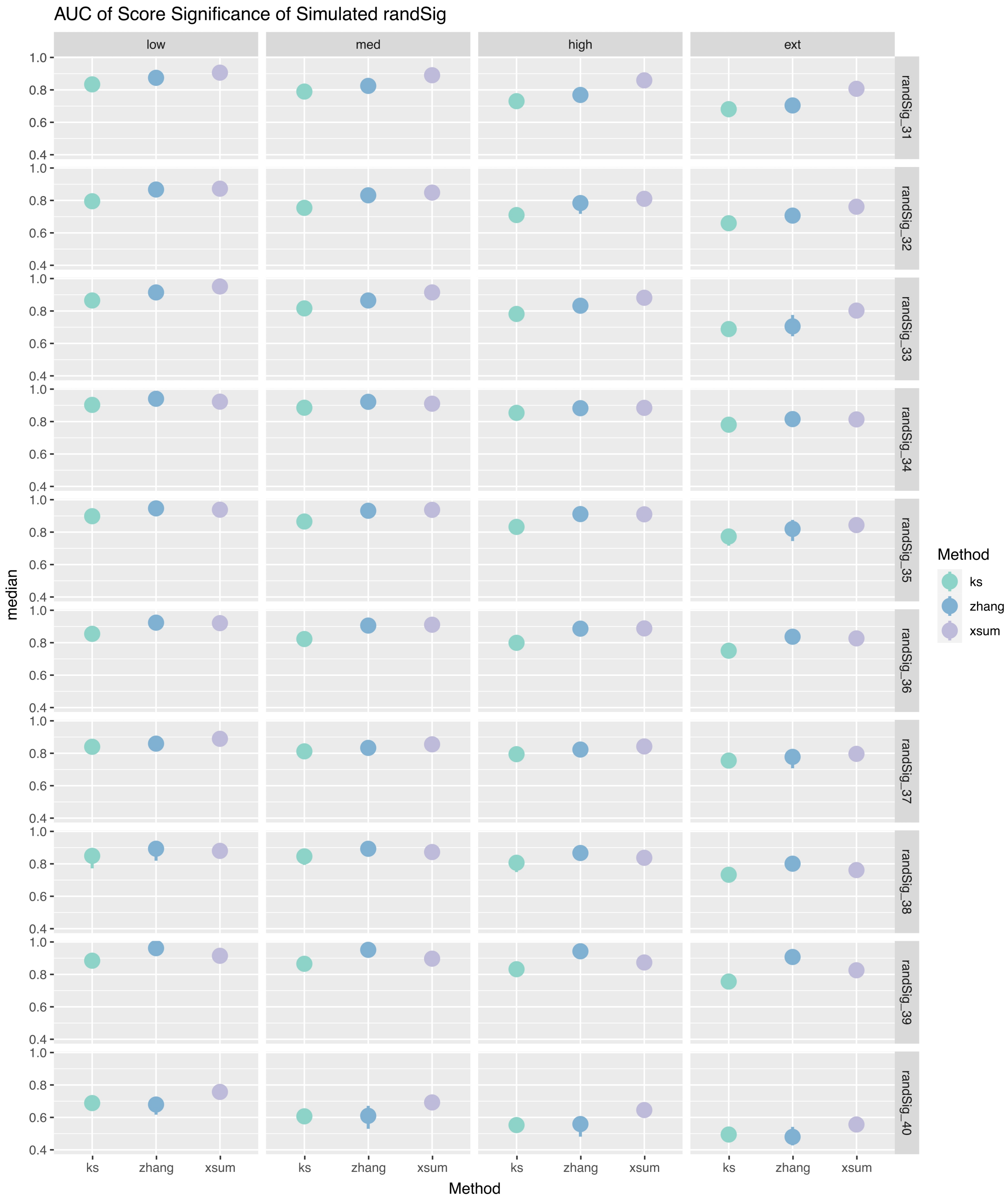

AUC of Score Significance of Simulated randSig
low
med
high
ext
randSig_31
randSig_32
randSig_33
randSig_34
randSig_35
randSig_36
randSig_37
randSig_38
randSig_39
randSig_40
median
1.0
0.8
0.6
0.4
ks
zhang
xsum
Method
Method
ks
zhang
xsum

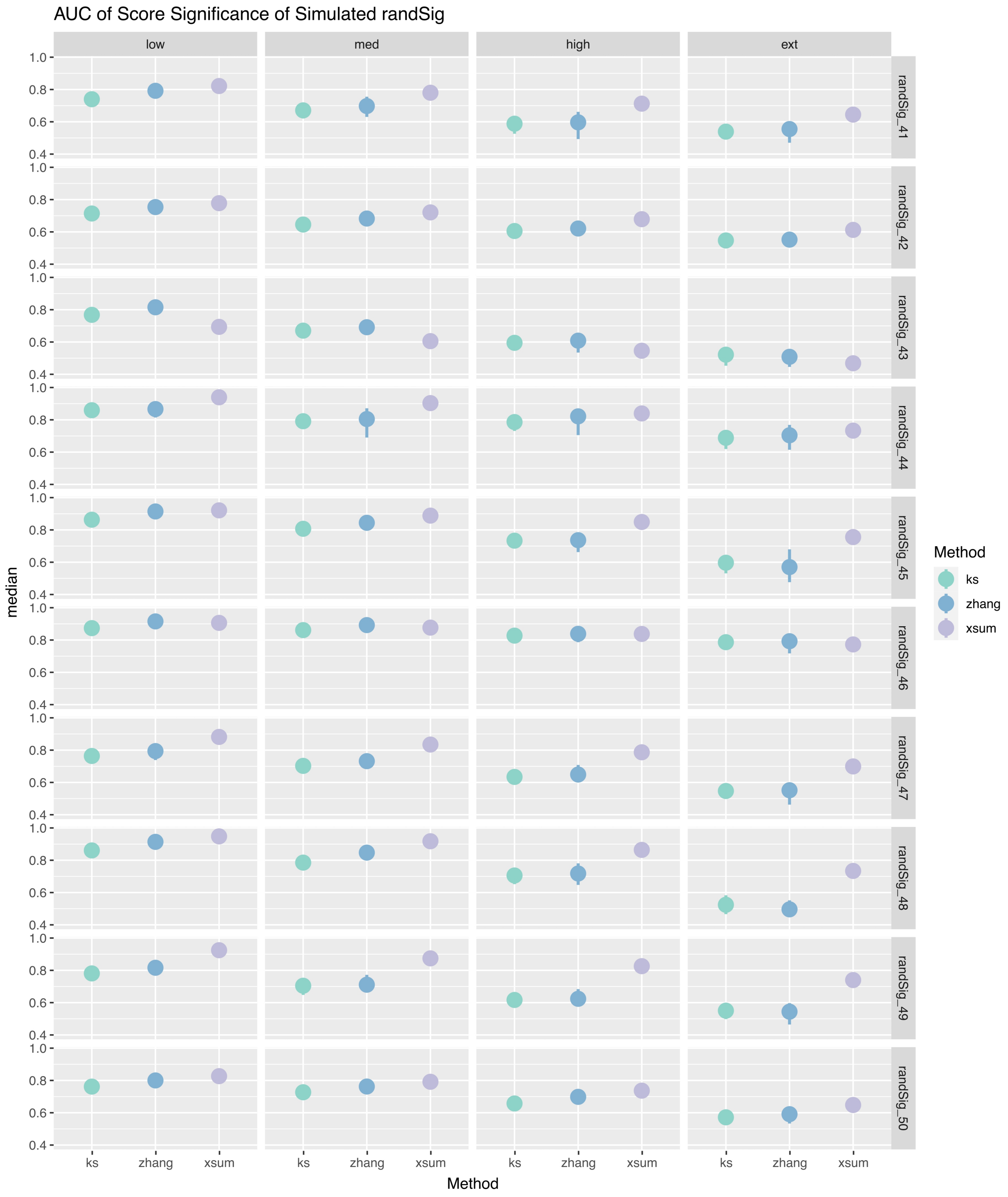


**Supplementary Figure 2C**. Point range plots of the median AUC (n=50) between the actual and predicted significance labels of randSig.

**Supplementary Table 1**. List of 50 randSigs that were selected for noise simulation in this study.

| RandSig Index | LINCS Signature ID |
|---|---|
| randSig_01 | AML001_CD34_6H:BRD-K49071277:10 |
| randSig_02 | LJP007_CD34_24H:D24 |
| randSig_03 | LJP007_CD34_24H:I23 |
| randSig_04 | LJP008_CD34_24H:J19 |
| randSig_05 | LJP009_CD34_24H:E15 |
| randSig_06 | LJP009_CD34_24H:E22 |
| randSig_07 | LJP009_CD34_24H:P02 |
| randSig_08 | LJP008_HEK293_24H:M15 |
| randSig_09 | LJP009_HEK293_24H:B12 |
| randSig_10 | REP.A013_HEK293_24H:N03 |
| randSig_11 | REP.A019_HEK293_24H:K22 |
| randSig_12 | REP.A024_HEK293_24H:K08 |
| randSig_13 | REP.B016_HEK293_24H:A15 |
| randSig_14 | REP.B023_HEK293_24H:E07 |
| randSig_15 | ASG003_MCF7_48H:O24 |

| | |
|---|---|
| randSig_16 | CPC011_MCF7_24H:BRD-K90789829-003-10-8:10 |
| randSig_17 | CPC017_MCF7_24H:BRD-A99177642-001-02-5:10 |
| randSig_18 | CPD001_MCF7_24H:BRD-K21401044-001-01-6:10 |
| randSig_19 | LJP001_MCF7_6H:BRD-K41895714-001-03-0:0.4 |
| randSig_20 | LJP009_MCF7_24H:M22 |
| randSig_21 | LKCP002_MCF7_24H:F09 |
| randSig_22 | RAD001_MCF7_24H:BRD-K15108141-003-01-3:0.0412 |
| randSig_23 | REP.A008_MCF7_24H:J23 |
| randSig_24 | REP.A025_MCF7_24H:E24 |
| randSig_25 | REP.A027_MCF7_24H:K07 |
| randSig_26 | REP.B026_MCF7_24H:K08 |
| randSig_27 | CPC012_NEU_24H:BRD-K74761218-001-03-5:10 |
| randSig_28 | CPC013_NEU_24H:BRD-K16478699-001-01-9:10 |
| randSig_29 | CPC014_NEU_24H:BRD-K06080977-001-13-3:10 |
| randSig_30 | CPC016_NEU_24H:BRD-A63836183-362-01-1:10 |
| randSig_31 | LJP007_NEU_24H:B08 |
| randSig_32 | LJP008_NEU_24H:B22 |

| | |
|---|---|
| randSig_33 | LJP008_NEU_24H:F23 |
| randSig_34 | LJP008_NEU_24H:O02 |
| randSig_35 | LJP009_NEU_24H:P21 |
| randSig_36 | NMH001_NEU_24H:BRD-K32814891-001-01-1:10 |
| randSig_37 | NMH001_NEU_6H:J24 |
| randSig_38 | NMH001_NEU_6H:O15 |
| randSig_39 | CEGS001_NPC.170_24H:C07 |
| randSig_40 | CEGS001_NPC.177_24H:J03 |
| randSig_41 | CEGS001_NPC.177_24H:L12 |
| randSig_42 | CEGS001_NPC.177_24H:L14 |
| randSig_43 | CEGS001_NPC.8330_24H:M13 |
| randSig_44 | LJP007_NPC.TAK_24H:I04 |
| randSig_45 | LJP008_NPC.CAS9_24H:G09 |
| randSig_46 | LJP009_NPC_24H:I16 |
| randSig_47 | LJP009_NPC.CAS9_24H:A17 |
| randSig_48 | LJP009_NPC.TAK_24H:J02 |
| randSig_49 | LJP009_NPC.TAK_24H:P21 |

| | |
|---|---|
| randSig_50 | NMH002_NPC_6H:BRD-K77830450-001-03-2:10 |
